# Endothelial ACKR1 expression regulates neutrophil infiltration and breast cancer metastatic engraftment in the lung metastatic niche

**DOI:** 10.64898/2026.03.15.711832

**Authors:** S. Tanner Roach, Qianxun Wang, Rishi Patel, Serena Thomas, Braulio Aguilar, Chinwe Ewenighi, Lauren Raasch, William A Muller, L.A. Naiche, Jan Kitajewski

## Abstract

The formation of the premetastatic niche prepares distant tissues for tumor cell engraftment. Endothelial cells are critical mediators of premetastatic niche formation, orchestrating extravasation of circulating tumor cells and critical pro-tumor immune cells, such as neutrophils. In mouse models of breast cancer, we show that primary tumors upregulate the non-signaling chemokine receptor ACKR1 in the endothelium of the lung premetastatic niche. ACKR1-expressing venules were found to be preferential sites of neutrophil and tumor cell localization within lung tissue. A newly generated conditional ACKR1 allele was used to show that endothelial-specific removal of ACKR1 expression significantly reduces metastatic engraftment in the lung. When ACKR1 is activated by tumor-secreted factors, endothelial ACKR1 functions to promote neutrophil recruitment within the lung parenchyma. We conclude that ACKR1 is a critical component of the endothelial response to tumors at the metastatic site of the lung, leading to neutrophil recruitment and promotion of tumor cell metastasis.

**SUMMARY:** Endothelial cells play critical roles in breast cancer metastasis. ACKR1 is upregulated in the endothelium of the lung metastatic niche in response to primary mammary tumors. Endothelial ACKR1 expression was found to promote neutrophil infiltration into the metastatic niche and support breast tumor cell metastasis to the lung.

## INTRODUCTION

Metastasis is responsible for the majority of breast cancer-related deaths^1^. The metastatic cascade is a multi-step process in which tumor cells detach from the primary tumor, intravasate into the circulation, and ultimately disseminate and colonize distant tissues. It is well established that distant tissues can be actively primed to support metastasis before tumor cells arrive, known as pre-metastatic niche formation^2-4^. The premetastatic niche is established via signals emanating from the primary tumor, which alter immune cell populations and endothelial function at the prospective metastatic site^2,3,5-7^. A growing body of evidence suggests that endothelial activation is an early event in premetastatic niche formation, creating an inflammatory microenvironment that promotes immune cell recruitment^8,9^. Tumor-secreted factors, including TNF-α, TGF-β, and VEGF-A, drive endothelial activation in the lung premetastatic niche prior to tumor cell arrival, leading to increased vascular permeability and adhesion molecule expression, which enhance tumor cell and immune cell infiltration^2,6,10^.

Neutrophils are among the first cells recruited to the premetastatic niche and have emerged as key regulators of metastasis^11-14^. Neutrophils are an abundant and heterogeneous innate immune cell type that function as first responders to infection or tissue injury. In the lung premetastatic niche, inflammatory signaling induces the expression of chemotactic signals, including CXCL1, CXCL2, and S100A8/A9, which recruit neutrophils to the metastatic niche^15,16^. Once recruited, neutrophils promote metastasis through several distinct mechanisms. Neutrophils can form neutrophil extracellular traps, which sequester circulating tumor cells and facilitate their colonization within the lung^17^. Neutrophils impair natural killer and T cell activity within the lung metastatic niche, creating an immunosuppressed microenvironment that enables tumor cells to evade immune surveillance^18-20^. Neutrophils secrete leukotrienes, which directly enhance the proliferation of metastasis-initiating cells^21^. Blocking neutrophil recruitment to the lung premetastatic niche has been shown to reduce tumor cell colonization and outgrowth in the lung, indicating an essential role in metastatic progression^15,21-23^. The ability of neutrophils to enhance metastatic progression suggests that elucidating the mechanisms of neutrophil trafficking to the premetastatic niche could enable therapeutic targeting to inhibit metastasis.

Atypical chemokine receptor 1 (ACKR1, also known as DARC or Duffy antigen) has emerged as a major regulator of inflammatory processes and neutrophil trafficking. ACKR1 is a seven-transmembrane receptor expressed primarily on red blood cells and on the venular endothelium of many organs, particularly in post-capillary venules and high endothelial venules of lymph nodes, which are preferential sites of extravasation for many circulating cell types^24-28^. ACKR1 is upregulated in endothelial cells of inflamed tissues, particularly in post-capillary venules and small veins that do not normally express this receptor under homeostatic conditions^29-35^.

Unlike classical chemokine receptors, ACKR1 does not trigger known intracellular signaling pathways, but instead binds and transports chemokines from the CC and CXC chemokine families, facilitating their retention and/or clearance^36-39^. ACKR1 is thought to increase chemokine concentrations at the luminal surface of inflamed endothelium via two mechanisms: ACKR1 can transport chemokines secreted by inflamed tissue from the abluminal to the luminal surface of endothelial cells^40,41^, or ACKR1 can immobilize chemokines in circulation or expressed by circulating cells, localizing them to the endothelial membrane^42^. Generation of a localized cytokine gradient by ACKR1 has been shown to promote neutrophil extravasation *in vitro* and *in vivo* ^32,40,42,43^. ACKR1 binds CXCL1 to enhance neutrophil adhesion to the endothelium and interacts with CXCL2 at endothelial cell junctions to promote neutrophil transendothelial migration^42,43^.

Changes in ACKR1 expression has been linked to clinical outcomes in breast cancer. In human breast cancer, low overall expression of ACKR1 in bulk RNA homogenate correlates with poor prognosis, but it is unclear which of the several cell types expressing ACKR1 are responsible for this effect^44^. Patients with low ACKR1 expression on red blood cells, known as the “Duffy null” blood group, show a higher incidence and mortality of triple-negative breast cancer (TNBC)^45,46^. In breast cancer tumor cells, higher expression of ACKR1 correlates with improved immune response in humans^44,46^, and reduced metastatic burden in mouse xenograft models^47,48^. However, the majority of ACKR1 protein is found on endothelial cells and red blood cells rather than tumor cells, suggesting that non-tumor cell sources of ACKR1 expression may underly the link to poor outcomes.

Although endothelial ACKR1 has been extensively studied in the context of inflammation and neutrophil recruitment, its contribution to cancer metastasis has not been directly examined. A recent study found that ACKR1 expression on endothelial cells of the primary tumor appears to correlate with increased cytotoxic immune infiltrate^49^. It is unknown whether ACKR1 expression in the lung endothelium is a feature of metastatic niche formation and how endothelial ACKR1 relates to neutrophil accumulation during metastatic progression.

In this study, we demonstrate that primary breast tumors or tumor conditioned media can induce ACKR1 expression in the endothelium of the lung pre-metastatic and metastatic niche, and that expression of endothelial ACKR1 is required for metastatic engraftment in the lung in both spontaneous and experimentally induced models of metastasis. ACKR1-expressing venules are a preferential site of neutrophil and tumor cell extravasation in the lung metastatic niche. We further show that endothelial ACKR1 regulates neutrophil recruitment to the lung metastatic niche. These findings identify a previously unrecognized tumor–endothelial crosstalk mechanism in which tumor-derived signals induce endothelial ACKR1 expression in the lung metastatic niche, resulting in both neutrophil accumulation and metastatic colonization. Targeting ACKR1 function may offer new therapeutic strategies for preventing metastatic disease.

## RESULTS

### ACKR1 is required to promote breast cancer metastasis to the lung

To determine the role of stromal ACKR1 in breast cancer growth and metastatic spread, we examined mammary tumor growth of orthotopically implanted tumor cells in mice with global deletion of *Ackr1* (Ackr1^Null^ mice)^50^. Tumor development was evaluated in immunocompetent mice implanted with different syngeneic mammary cell lines: two independently derived MMTV-PyMT mammary carcinoma lines, PY8119 and AT3, and a lung-tropic metastatic mouse luminal B cancer cell line, E0771.LMB (LMB)^51-54^. LMB cells have low endogenous ACKR1 expression (Fig. S1A). We orthotopically implanted 2x10^5^ LMB or PY8119 cancer cells into the fourth mammary fat pad of immunocompetent Ackr1^Null^ or wildtype C57BL6/J mice (WT). Minimal ACKR1 expression was observed in tumor cells or tumor endothelium within the primary tumor, however, some vessels around the periphery of the tumor expressed ACKR1 in wild-type mice (Fig. S1B). LMB and PY8119 tumors implanted in Ackr1^Null^ animals did not grow at a significantly different rate in WT or Ackr1^Null^ animals, suggesting that the loss of host tissue ACKR1 does not have a major effect on primary mammary tumor growth (Fig. S1C-E).

Metastasis to the lung is common in breast cancer and one of the most significant predictors of poor outcomes^55,56^. We therefore examined the lungs of mammary tumor-bearing mice. PY81119 cells do not spontaneously metastasize, so lungs from PY8119-bearing mice were not examined. We found that Ackr1^Null^ mice with orthotopic LMB cell implantation contained few or no detectable tumor cells in the lungs, indicating that loss of ACKR1 from host tissue significantly reduced metastasis to the lungs (Fig.1A-B, Fig. S1F). These results indicate that host tissue expression of ACKR1 is required for breast cancer metastasis to the lung.

**Figure 1.**
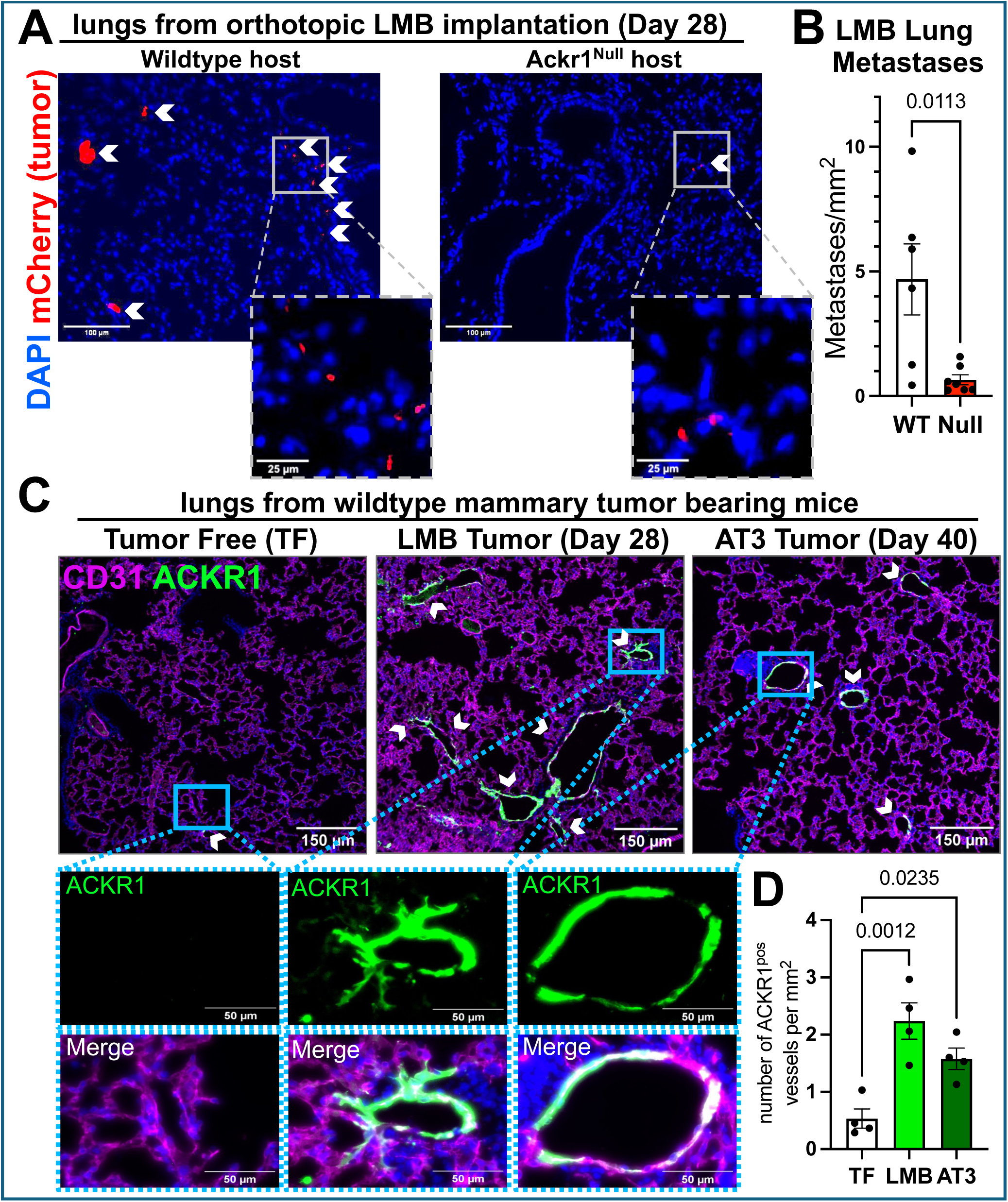
Host ACKR1 expression is required for breast cancer metastasis to the lung. **(A)** Representative images of spontaneous lung metastases from orthotopic E0771.LMB (LMB) mammary tumors implanted into WT and Ackr1^Null^ mice harvested at humane endpoint (2 cm tumor diameter), Day 28 after tumor injection. Tumor cells are detected with mCherry (red) with individual tumor cells (white arrowheads) indicated in higher magnification inset. **(B)** Quantification of lung metastases in WT (n = 6) and Ackr1^Null^ (n = 7) mice. Error bars indicate s.e.m. in all figures. Significance determined by unpaired t-test. Each data point represents a single mouse, in which a minimum of three sections at different depths within the lung have been analyzed and averaged. **(C)** Representative lung sections from tumor-free (TF) mice and mice bearing LMB and AT3 orthotopic mammary tumors harvested at humane endpoints (Day 28 and Day 40, respectively). Immunofluorescent (IF) staining of endothelial cells with CD31 (magenta), ACKR1 (green) and DAPI (blue). ACKR1-expressing vessels indicated with white arrowheads. **(D)** Quantification of ACKR1-positive vessels per mm^2^ in TF, LMB-bearing, and AT3-bearing mouse lungs. Significance determined by one-way ANOVA with Sidak’s multiple comparisons test.

To investigate the role of ACKR1 in lung metastasis, we examined ACKR1 expression in the lung. We found that tumor free (TF) WT mice showed little or no ACKR1 expression in the lung endothelium or parenchyma (Fig. 1C). Mice bearing orthotopic LMB tumors exhibited strong endothelial ACKR1 expression, with a ∼4-fold increase in the number of ACKR1-positive vessels at humane endpoint (i.e. 2 cm tumors, which occurred on day 28 after tumor implantation for LMB cells and day 40 for AT3 cells) (Fig. 1C-D). Similarly, orthotopic implantation of the metastatic AT3 cell line significantly increased the number of ACKR1-positive vessels in pulmonary endothelium at humane endpoint (Fig 1C-D). These results indicate that ACKR1 expression is upregulated in the lung endothelium of mice with primary tumors and that the ability of tissue to express ACKR1, possibly in the vasculature, promotes breast cancer metastasis to the lung.

### Endothelial ACKR1 is upregulated in the lung pre-metastatic niche of tumor-bearing mice

We sought to determine the timing of ACKR1 upregulation in the lung endothelium of tumor bearing mice; assessing if ACKR1 regulation occurs during premetastatic niche formation or after metastatic engraftment. We longitudinally examined lung endothelial ACKR1 expression at 3, 7, 13, and 28 days after orthotopic implantation of LMB cells into WT mice (Fig 2A, D and Fig. S2A). Negligible numbers of tumor cells were detected in the lungs prior to Day 28 (Fig. 2C). We found significant upregulation of endothelial ACKR1 expression in the lung vasculature of tumor-bearing mice starting at 13 days, indicating that ACKR1 upregulation precedes tumor cell seeding (Fig. 2A, D).

**Figure 2.**
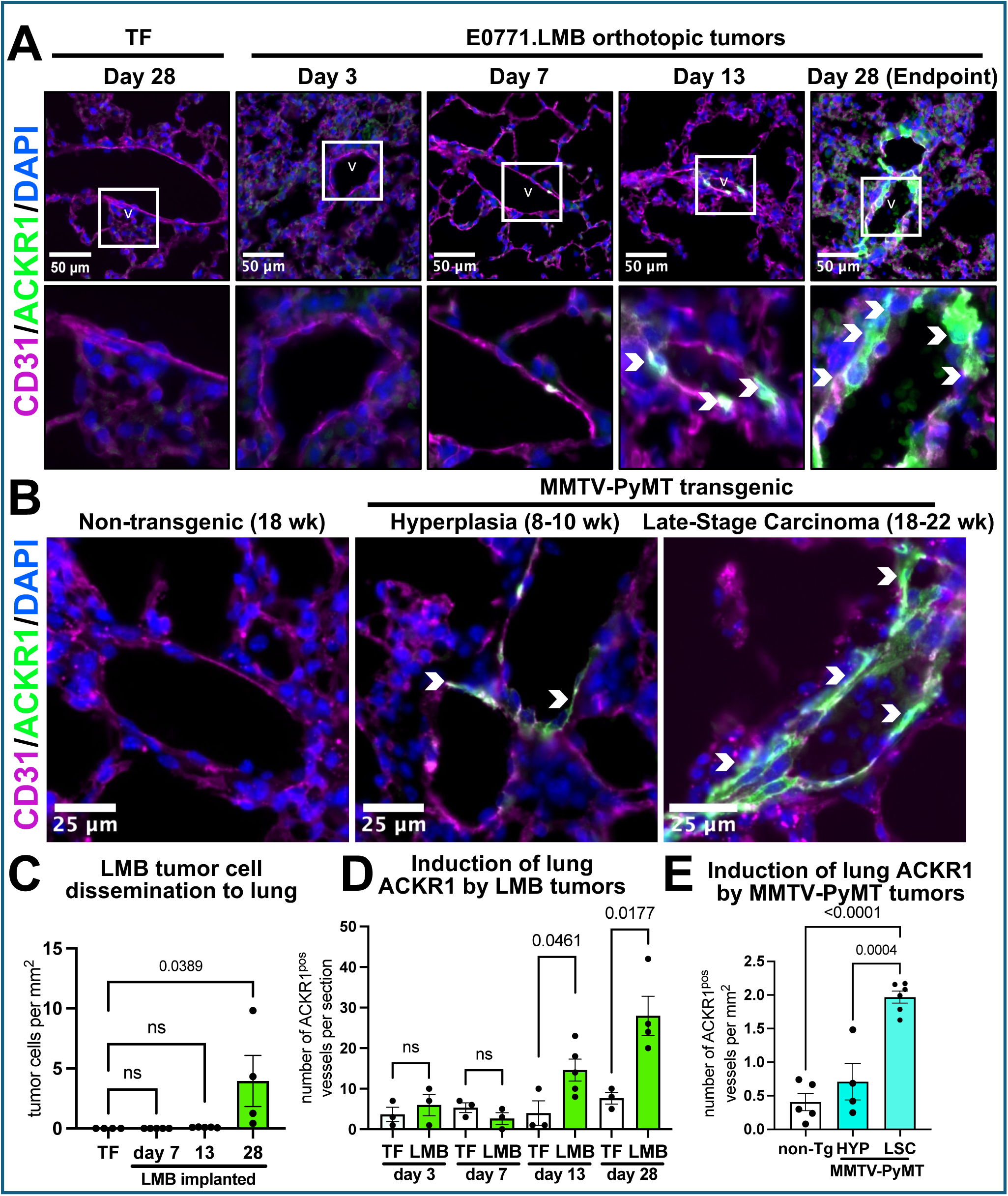
Endothelial ACKR1 is activated in the lung pre-metastatic niche. **(A)** Representative IF images of lungs from C57BL6/J wildtype mice at 3, 7, 13, and 28 days after orthotopic injection with LMB cells or tumor-free (TF) PBS mock injection. CD31 = magenta, ACKR1 = green, DAPI = blue in all panels. ACKR1-positive endothelial cells indicated by white arrowheads. Fainter ACKR1 staining can also be seen in red blood cells. **(B)** Representative IF images of lungs from non-transgenic or MMTV-PyMT transgenic C57BL6/J mice at 8-10 weeks (hyperplasia stage) and 18-22 weeks (late stage carcinoma). **(C)** Number of disseminated LMB tumor cells per mm^2^ in lungs at indicated timepoints after tumor implantation. One-way ANOVA with Sidak’s multiple comparisons test. **(D)** Average number of ACKR1-positive vessels per lung section at indicated timepoints after tumor implantation. Unpaired t-test. **(E)** Quantification of ACKR1-positive vessels per mm^2^ in lungs of non-transgenic (18 week old), MMTV-PyMT transgenic at hyperplasia stage (8-10 week old, HYP) and MMTV-PyMT transgenic at late stage carcinoma (18-22 week old, LSC) mice. One-way ANOVA with Sidak’s multiple comparisons test. For all panels, each data point represents a single mouse, in which a minimum of three sections at different depths within the lung have been analyzed and averaged.

To evaluate the tumor-derived regulation of lung endothelial ACKR1 in a spontaneous mammary tumorigenesis setting that closely mimics human disease, we examined lungs from mice carrying the mouse mammary tumor virus-polyoma middle T antigen (MMTV-PyMT) transgene, which recapitulates the progression of naturally arising breast cancer, including hyperplasia, early carcinoma, late stage carcinoma, and metastatic disease^57-59^. In this model, mice generally start to show metastasis starting at ∼12 weeks old^60,61^. We found that MMTV-PyMT mice with mammary hyperplasia (8-10 week old, HYP) showed detectable upregulation of ACKR1 in the lungs. Late-stage carcinoma MMTV-PyMT mice (18-22 weeks old, LSC) had significantly increased lung endothelial ACKR1 expression compared to non-transgenic mice (non-Tg) (Fig. 2B, E, and Fig. S2B). This timing indicates that ACKR1 is upregulated in lung endothelium prior to or concurrent metastatic seeding.

### Mammary tumor cell-secreted factors are sufficient to upregulate endothelial ACKR1

Prior work has shown that ACKR1 can be upregulated in endothelial cells by secreted inflammatory factors [i.e. TNF or lipopolysaccharide (LPS)] or by direct cell-cell contact with neutrophils^62^. We posited that lung endothelial Ackr1 could be upregulated by factors secreted from the primary tumor or could require direct contact with disseminated tumor cells in circulation. To determine whether factors secreted from the primary tumor can induce ACKR1 expression, we harvested supernatant media that had been incubated with mammary tumor cells for 48 hours (tumor conditioned media, TCM) or had not been incubated with cells (tumor free media, TFM) and snap frozen (Fig 3A). We intraperitoneally injected 300 μl TFM or TCM from LMB cells (TCM-LMB) into WT mice daily for 7 days. We observed a significant increase in ACKR1-positive vessels in the lungs of TCM-LMB treated mice at the day 7 timepoint (Fig. 3B-C and Fig. S2C). Thus, tumor-secreted factors can induce ACKR1 expression in the lung endothelium without requiring metastatic cell contact, consistent with a role for ACKR1 in the lung premetastatic niche.

**Figure 3.**
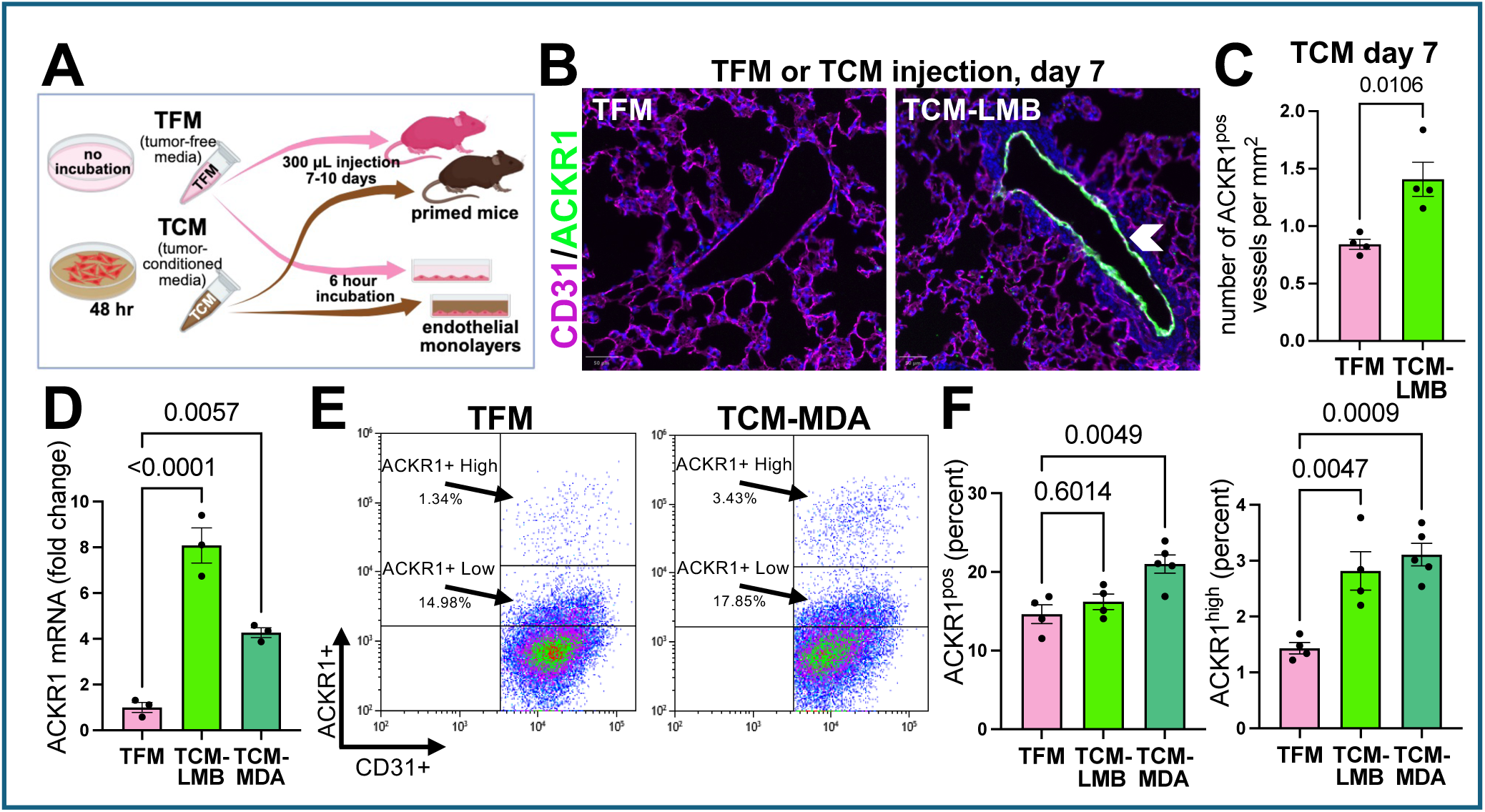
Secreted factors from tumor cells are sufficient to induce endothelial ACKR1. **(A)** Schematic of tumor-conditioned media production and administration. **(B)** Representative IF images of lungs from C57BLJ/6 wildtype mice treated daily for 7 days with 300µl tumor-free media (TFM) or media conditioned by incubation with LMB tumor cells (TCM-LMB) (CD31 = magenta, ACKR1 = green, DAPI = blue); ACKR1-expressing venule is indicated by white arrowhead. **(C)** Quantification of ACKR1-positive vessels per mm^2^ in lungs of mice treated with TFM or TCM for 7 days. Unpaired t-test. **(D)** qPCR of ACKR1 mRNA expression in HUVEC monolayers treated with TFM, MDA-MB-231-conditioned media (TCM-MDA), or TCM-LMB. One-way ANOVA with Sidak’s multiple comparisons test. **(E-F)** Flow cytometry analysis of cell surface ACKR1 expression in HUVEC treated with TFM or TCM-MDA for 24 hours. ACKR1^pos^ or ACKR1^high^ numbers expressed as a percentage of CD31^pos^ cells. One-way ANOVA with Sidak’s multiple comparisons test.

We tested whether induction of ACKR1 was a direct effect of tumor-secreted factors on endothelium, as opposed to requiring other cell types (i.e. immune cells) for this induction. We generated endothelial EGM2 media conditioned by LMB cells (TCM-LMB) or from the human metastatic breast cancer cell line MDA-MB-231 (TCM-MDA)^63^. We treated HUVEC monolayers with TFM, TCM-LMB, and TCM-MDA. After 6 hours of treatment, both LMB and MDA-conditioned media significantly increased ACKR1 mRNA expression compared to TFM (Fig. 3D). To determine whether this resulted in an increase in endothelial cell surface ACKR1 expression, we examined ACKR1 expression by flow cytometry after 24 hours of treatment. We found a significant increase in both the overall percentage of ACKR1^pos^ cells and the percentage of ACKR1^HI^ cells in TCM-MDA treated HUVECs (Fig. 3E-F). These results demonstrate that the endothelial cell ACKR1 upregulation in a mouse mammary tumor model can be replicated by secreted factors from human breast cancer cells interacting with human endothelial cells.

### Metastatic tumor cells and neutrophils preferentially localize near endothelial ACKR1-expressing pulmonary venules in tumor bearing mice

During metastasis, tumor cells extravasate from the vasculature into the surrounding parenchyma. Formation of the premetastatic and metastatic niche requires extravasation of neutrophils, which secrete factors that support tumor cell survival and engraftment^11,14^. In most tissues, extravasation is thought to occur primarily in post-capillary venules and small veins, although recent data suggests that neutrophils can also extravasate from capillaries in the lung^64-66^. We examined ACKR1 expression in day 28 LMB tumor-bearing mouse lungs stained for the venous endothelial marker Ephrin Receptor B4 (EphB4). We show that all endothelial ACKR1 expression localizes to EphB4-positive vessels (Fig. 4A and Fig. S3A-B), indicating that endothelial ACKR1 upregulation specifically occurs in pulmonary venules. Only a subset of EphB4-positive venules expressed ACKR1, but the percentage of venules that expressed ACKR1 was consistent between animals (22.7% ± 1.7 s.e.m., n = 4 mice) (Fig. S3B).

**Figure 4.**
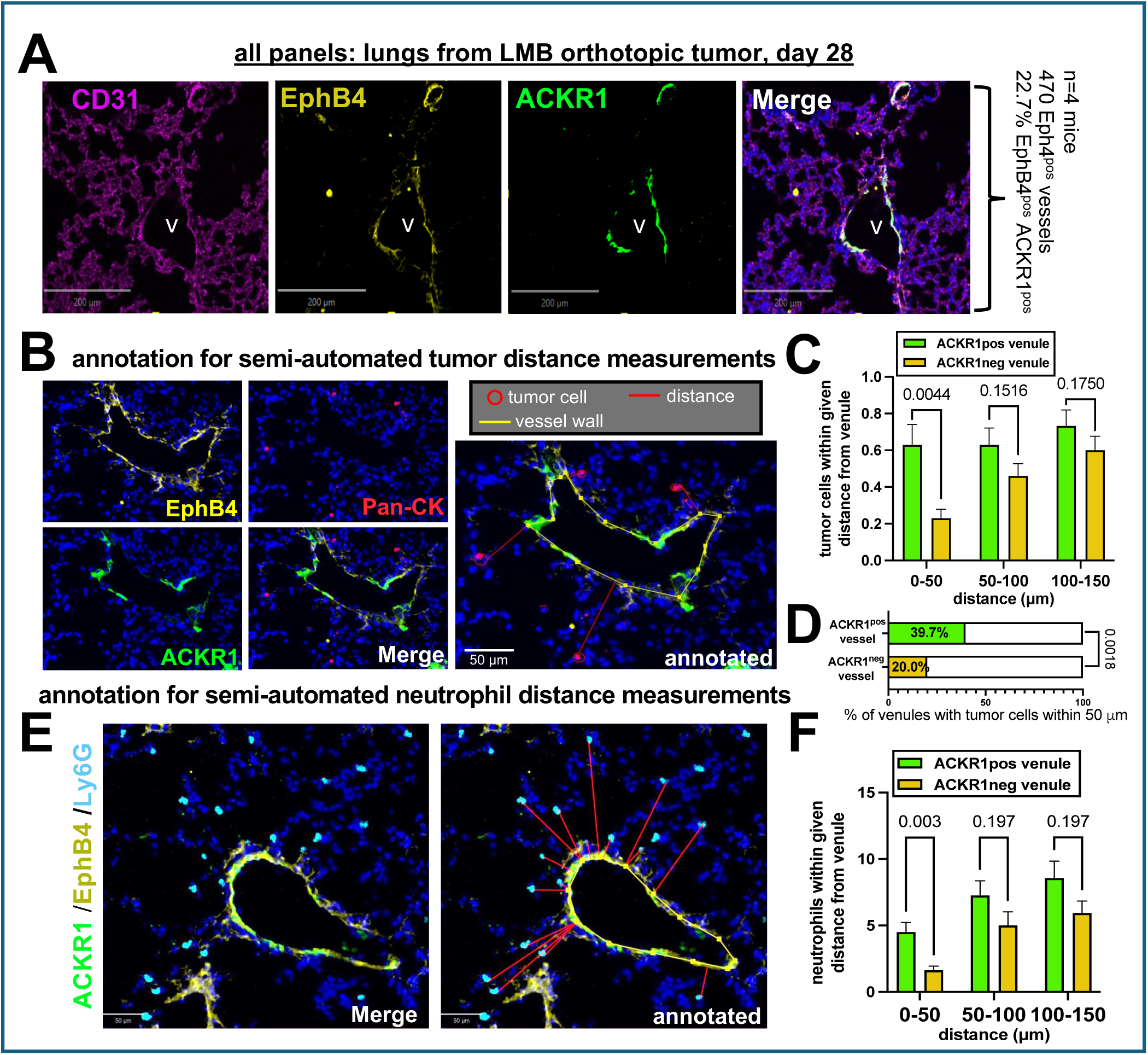
Disseminated tumor cells and neutrophils preferentially localize near ACKR1-expressing pulmonary venules. **(A)** Representative image of ACKR1-positive lung vessel from LMB-bearing mouse 28 days after implantation, showing colocalization of ACKR1 with venous endothelial marker EphB4. CD31 = magenta, EphB4 = yellow, ACKR1 = green. All ACKR1^pos^ vessels also expressed EphB4. 22.7±1.4% of EphB4^pos^ venules expressed ACKR1. Quantitation shown in Supplemental Fig. S3B. **(B)** Representative annotated IF image for QuPath-based proximity analysis of tumor cell distance to nearest venule (EphB4 = yellow, Pan-CK = tumor cell marker pancytokeratin in red, ACKR1 = green, and DAPI = blue). For QuPath annotations, yellow line = vessel border, red circle = disseminated tumor cell (DTC), red line = shortest distance between DTC and nearest venule border. **(C)** Quantification of the number of DTCs within indicated distance from the border of ACKR1-positive or ACKR1-negative venules in Day 28 LMB-bearing mouse lungs (n = 116 ACKR1^pos^ venules, n = 100 ACKR1^neg^ venules, across 4 individual mice). Unpaired t-tests. Summed data shown here for clarity, individual data points shown in Supplemental Fig. S3C. **(D)** Percentage of ACKR1-positive and ACKR1-negative venules with at least one DTC within 50 µm of the venule border. Chi-squared test. **(E)** Representative IF images for QuPath proximity analysis of neutrophil distance to nearest venule, as described in panel B except Ly6G = neutrophil marker (cyan) and red lines indicate shortest distance between neutrophil and nearest venule border. **(F)** Quantification of the number of neutrophils within indicated distance from the border of ACKR1-positive or ACKR1-negative venules in Day 28 LMB-bearing mouse lungs (n = 16 ACKR1^pos^ venules, n = 16 ACKR1^neg^ venules, across 3 individual mice). Unpaired t-tests. Summed data shown here for clarity, individual data points shown in Supplemental Fig. S3D.

In inflammatory settings, endothelial ACKR1 expression has been shown to promote the extravasation of leukocytes, particularly neutrophils^26,27,40,42^. Extravasation of circulating tumor cells into the premetastatic niche frequently mirrors the mechanisms by which leukocytes exit the bloodstream^67^. We therefore tested whether metastatic tumor cells and neutrophils preferentially localize near ACKR1-positive vessels, an indicator that these vessels act as sites of extravasation for each respective cell type.

To determine if disseminated tumor cells localized near ACKR1-positive vessels, we stained sections from Day 28 LMB tumor-bearing mouse lungs with pan-cytokeratin (Pan-CK, tumor cell marker), ACKR1, and EphB4. Using blinded vessel annotation and a custom QuPath macro, we measured the distance of disseminated tumor cells to the nearest pulmonary venule (Fig. 4B). We found a ∼2.8 fold enrichment of tumor cells within 50 μm from ACKR1-positive venules as compared to ACKR1-negative venules (Fig. 4C and Fig. S3C). Significantly more ACKR1-positive venules had at least one tumor cell within 50µm of the vessel border than ACKR1-negative venules (Fig. 4D).

Similar analysis using Ly6G to identify neutrophils showed significantly higher numbers of neutrophils within 50 μm of ACKR1-positive venules compared to ACKR1-negative venules (Fig. 4E-F and Fig. S3D). These data show that tumor cells and neutrophils are enriched near the subset of venules that express ACKR1 in the lung metastatic niche, indicating that ACKR1-positive venules are the likely site of extravasation for both tumor cells and neutrophils.

### Generation of an endothelial-cell specific ACKR1 conditional knockout mouse model

To evaluate cell-type specific roles of ACKR1 in breast cancer lung metastasis, we generated a conditional *Ackr1* knockout mouse (Ackr1^flox^). Using CRISPR-mediated targeted mutagenesis, we introduced *loxP* sites around exon 2 of the *Ackr1* allele, which encodes nearly the entire ACKR1 protein (Fig. 5A). Human mutations in a GATA1/TAL1 binding site in the ACKR1 promoter region disrupt ACKR1 expression in red blood cells (RBC), known as the Duffy null phenotype, so we positioned the loxP sites to avoid murine consensus GATA1/TAL1 binding sites (Fig. 5A). Based upon our discovery of endothelial ACKR1 expression in the metastatic niche, we used the *Ackr1^flox^* allele to evaluate endothelial function of ACKR1 in metastatic spread. We crossed *Ackr1^flox^*mice with the endothelial-specific *Cdh5CreERT2* line^68^, producing *Ackr1^flox/flox^;Cdh5CreERT2* mice, referred to as Ackr1^ECKO^, and *Ackr1^wt/wt^;Cdh5CreERT2* mice, referred to as Controls. For all experiments, endothelial-specific loss of ACKR1 was induced by treating Ackr1^ECKO^ and Control mice with 2mg/day tamoxifen daily for 5 days via oral gavage starting at 4-5 weeks old.

**Figure 5.**
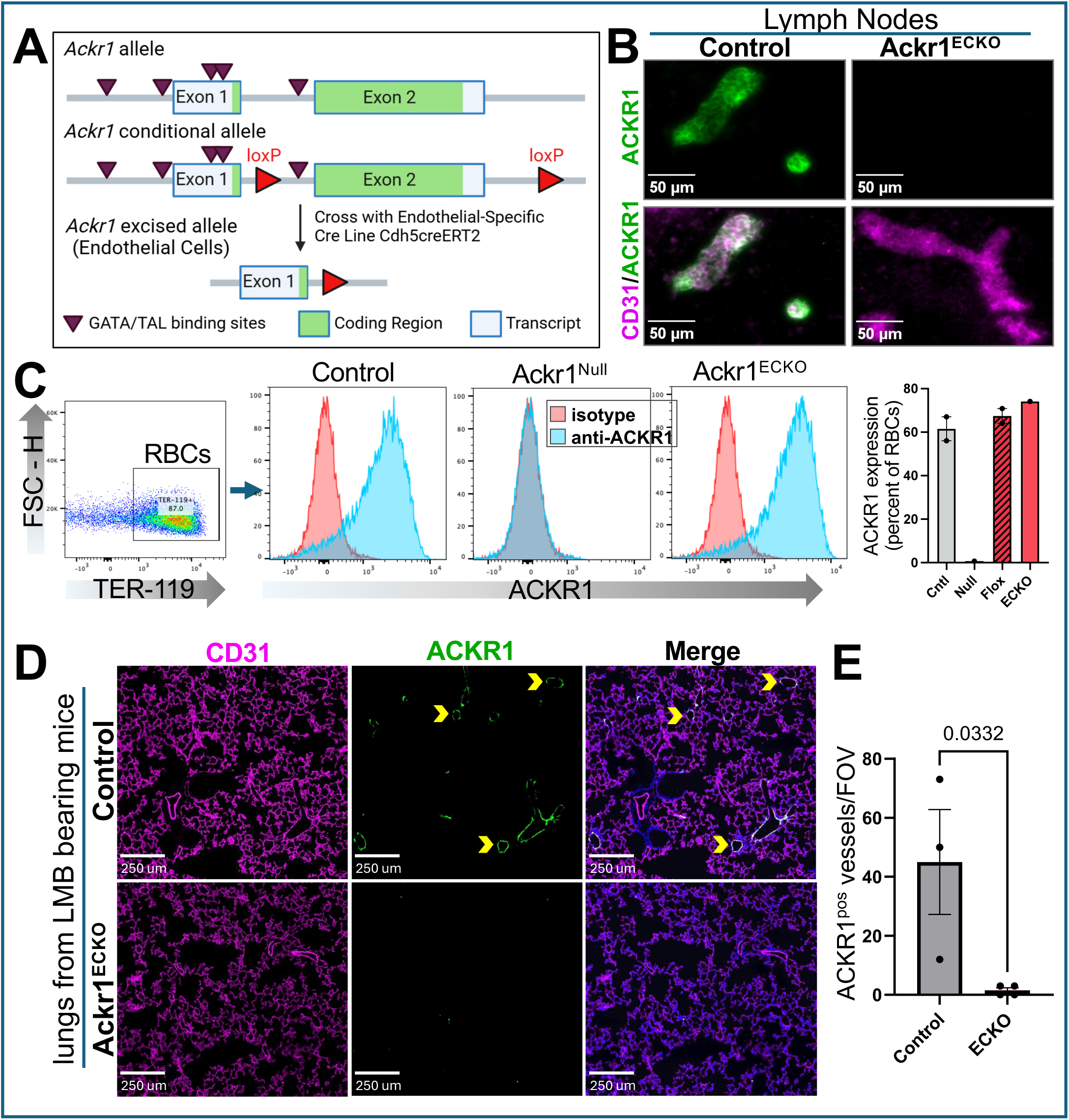
Generation of an endothelial cell-specific *Ackr1* conditional knockout mouse model. **(A)** Schematic of the *Ackr1* floxed allele and its excision in endothelial cells following tamoxifen-induced *Cdh5CreERT2* recombination to generate the Ackr1^ECKO^ mouse model. Consensus GATA/TAL binding sequences (black triangles) were not disrupted when inserting the loxP sites (red triangles). **(B)** Representative IF images (n=2 mice per genotype) of peripheral lymph nodes, which constitutively express ACKR1 in endothelial cells, confirming the loss of ACKR1 protein in Ackr1^ECKO^ mice (CD31 = magenta, ACKR1 = green). **(C)** Flow cytometry analysis of peripheral blood confirms retention of ACKR1 expression in red blood cells of Ackr1^ECKO^ mice. TER-119 = red blood cell marker, anti-ACKR1 = blue peaks, isotype control = red peaks. Control = *Cdh5CreERT2;Ackr1^WT^*, Null = Ackr1^Null^ (negative control), Flox = *Ackr1^flox/flox^* (no cre control), ECKO = *Cdh5CreERT2;Ackr1^flox/flox^* (experimental). **(D)** Representative images of Day 28 LMB mammary tumor-bearing Control and Ackr1^ECKO^ mouse lungs confirming loss of endothelial ACKR1 in Ackr1^ECKO^ mouse lungs after induction by primary mammary tumors (CD31 = magenta, ACKR1 = green). **(E)** Quantification of loss of ACKR1-positive vessels in Day 28 LMB tumor-bearing Control (n=3) and Ackr1^ECKO^ (n=4) mouse lungs. Unpaired t-test.

To validate endothelial-specific ACKR1 deletion, we examined high endothelial venules (HEVs) in peripheral lymph nodes, where ACKR1 is constitutively expressed^27^. ACKR1 expression was lost in Ackr1^ECKO^ mice HEVs (Fig. 5B). To confirm RBC ACKR1 expression was preserved, we stained peripheral blood from Control, Ackr1^ECKO^, and Ackr1^Null^ mice for ACKR1 and TER-119 (RBC marker) and analyzed expression via flow cytometry. Ackr1^ECKO^ mice showed comparable RBC ACKR1 expression as Control mice, whereas Ackr1^Null^ mice RBCs lacked ACKR1 expression (Fig. 5C). To test whether ACKR1 was efficiently excised in lung endothelium, we examined ACKR1 expression in LMB tumor bearing mice 28 days post implantation. Ackr1^ECKO^ mice showed a negligible number of ACKR1-positive vessels (Fig 5D-E). We therefore confirm the successful generation of an Ackr1^ECKO^ mouse model that preserves RBC ACKR1 expression while selectively ablating ACKR1 in endothelial cells.

### Endothelial ACKR1 promotes mammary tumor metastasis to the lung

Having established reduced lung metastasis in Ackr1^Null^ mice, we used the Ackr1^ECKO^ mouse model to evaluate whether endothelial ACKR1 is required for metastatic colonization in the lung. We tested the ability of breast cancer cells to engraft in the lung metastatic niche in Ackr1^ECKO^ mice. To avoid potential confounding factors of ACKR1 ablation in the primary tumor, we employed an experimentally induced metastasis model in which tumor cells are injected directly into the bloodstream and allowed to disseminate. We recapitulated primary tumor induction of lung endothelial ACKR1 by administering 300 μl TCM-LMB or TCM-AT3 to Control and Ackr1^ECKO^ mice for 8 days (called “priming”, Fig. 6A). We then injected fluorescently labeled LMB or AT3 mammary tumor cells, respectively, via the tail vein and allowed them to circulate and establish metastatic foci for two weeks (Fig. 6A). Lungs were examined for mCherry-positive metastases via whole mount fluorescence. Ackr1^ECKO^ mice showed a significant reduction in lung metastases of both LMB and AT3 cells (Fig. 6B-F). In the LMB model, the majority of Ackr1^ECKO^ animals showed no metastatic nodules (Fig. 6D). Collectively, these data demonstrate that lung endothelial ACKR1 is required to promote breast cancer cell engraftment and metastatic growth in the lung.

**Figure 6.**
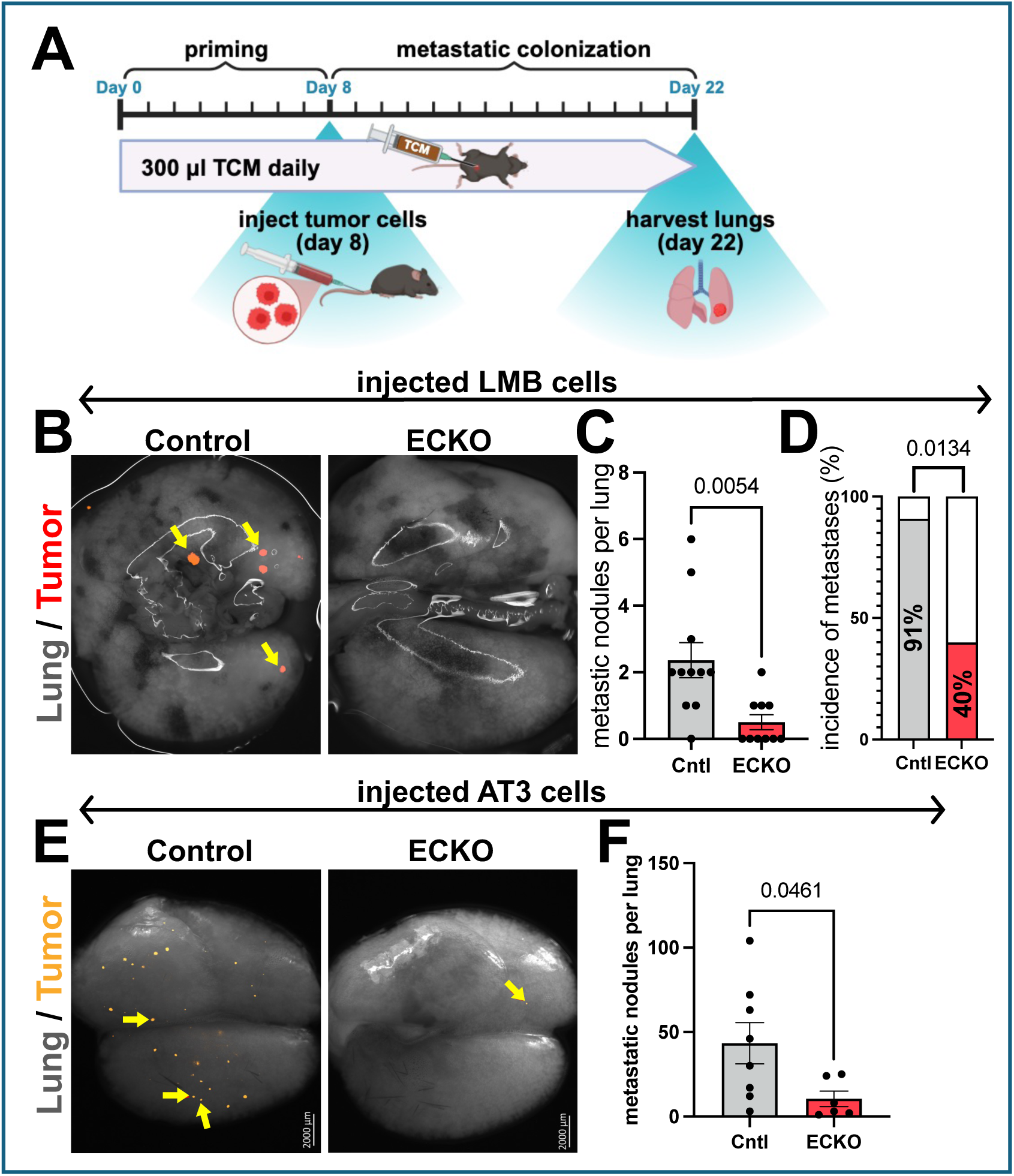
Endothelial ACKR1 promotes breast cancer metastasis to the lung. **(A)** Schematic of experimental lung metastasis protocol. 300 µl TCM-LMB or TCM-AT3 was administered daily via IP injection, then the corresponding tumor cells injected on day 8 via tail vein injection. Lungs were harvested 2 weeks after tumor cell injection (day 22). **(B)** Representative lung images from Control and Ackr1^ECKO^ mice following injection of LMB tumor cells (brightfield lung image = gray, metastatic LMB cells = mCherry in red/orange, yellow arrowheads = metastatic nodules). **(C)** Quantification of LMB lung metastases in Control (n = 11) versus Ackr1^ECKO^ (n = 10) mice. Unpaired t-test. **(D)** Percentage of mice which showed evidence of any metastatic nodules (incidence). Chi-squared test. **(E)** Representative lung images from Control and Ackr1^ECKO^ mice following injection of AT3 tumor cells (Colors same as panel B). **(F)** Quantification of AT3 lung metastases in Control (n = 8) versus Ackr1^ECKO^ (n = 6) mice. Unpaired t-test.

### Endothelial ACKR1 is not required for tumor cell extravasation in the lung

To examine the mechanism by which ACKR1 promotes lung metastasis, we tested whether endothelial ACKR1 regulates tumor cell recruitment and extravasation during lung metastasis. We primed mice with TCM-LMB to induce endothelial ACKR1, as described above, or with TFM to represent non-induced baseline conditions. Following 9 days of either TCM or TFM injections, mice received intravenous injections of fluorescently labeled LMB cells, which were allowed to disseminate for 24 hours. To distinguish cells remaining in circulation from extravasated cells, we administered anti-MHCI antibody intravenously three minutes prior to euthanasia to label all cells that remained in circulation (Figure 7A)^69^. Lungs were disassociated and analyzed via flow cytometry. We confirmed that all tumor cells that remained in the intravascular space were labeled by anti-MHCI by collecting whole blood from mice, using the mCherry signal to detect tumor cells. 99% of circulating tumor cells were stained positively for MHCI (Fig. S4A). Further, we confirmed that the intravenously injected antibody does not leak into the lung parenchyma, even under inflammatory conditions (Fig. S4B). We found no significant difference between Control and Ackr1^ECKO^ mice in either the total number of tumor cells present in the lungs or in the proportion of intravascular to extravasated tumor cells in either the TCM- and TFM-treated conditions (Fig 7B-C). These results indicate that in the LMB model of metastasis, endothelial ACKR1 does not directly regulate tumor cell recruitment to the lung or extravasation from the bloodstream into the lung parenchyma.

**Figure 7.**
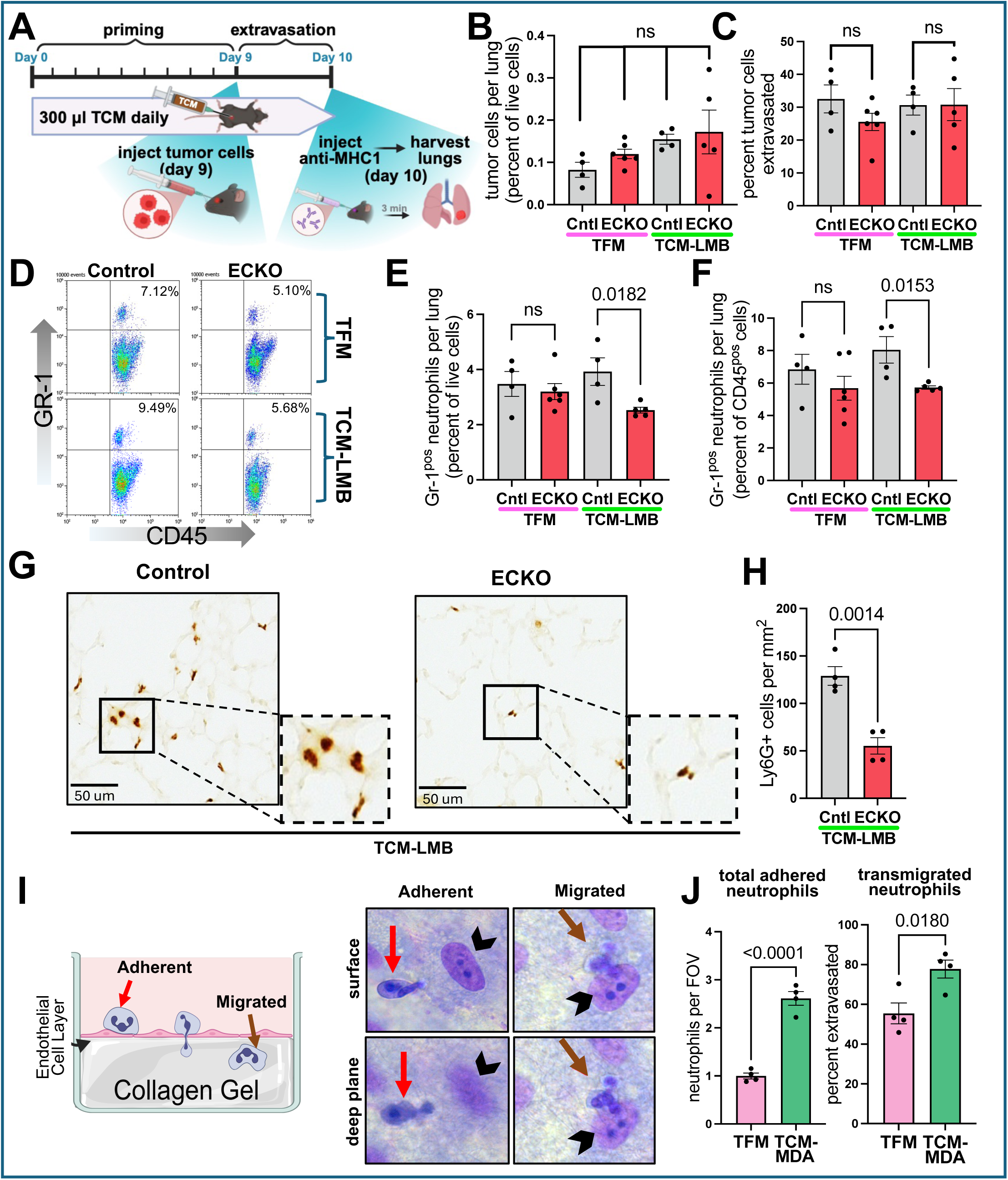
Endothelial ACKR1 regulates neutrophil recruitment to the lung metastatic niche. **(A)** Schematic of extravasation assessment protocol. 300 µl TFM or TCM-LMB was administered daily via IP injection, then LMB cells were administered on day 9 via retroorbital injection. 24 hours after tumor cell injection, anti-MHCI (H-2kB) antibody was retroorbitally injected to label intravascular tumor cells and allowed to circulate for 3 minutes before sacrifice. **(B)** Flow cytometry quantifying total tumor (RFP^pos^) cells in the lung. Each dot represents one mouse. One-way ANOVA with Tukey’s multiple comparisons. The same mice are represented in panels B-F. **(C)** Percentage of extravasated (RFP^pos^, MHC1^neg^) tumor cells in the lung. Unpaired t-tests. **(D)** Representative gating strategy for neutrophils (CD45+/Gr-1+). **(E)** Quantitation of neutrophil infiltrate, expressed as percentage of total lung cells. Unpaired t-tests. **(F)** Contribution of neutrophils to the total immune cell infiltrate (CD45+). Unpaired t-tests. **(G)** Representative immunohistochemistry images of lung neutrophils (Ly6G, brown). **(H)** Neutrophils per mm^2^ in lung tissue sections. Unpaired t-test. **(I)** Schematic illustrating the design of neutrophil transmigration assay. HUVEC were grown to confluence onto a collagen gel and cultured for 24 hours in TFM or MDA-TCM. HUVEC were either left unstimulated or activated with TNFα for 4 hours. Freshly harvested neutrophils (PMNs) were seeded on top and allowed to migrate for 6 minutes. Cultures were rinsed with 1mM EDTA to remove loosely adherent cells, then fixed and stained. In Z-stack images, neutrophils that were in focus in the same plane as endothelial cells (black arrowheads) were scored as surface adherent (red arrows) and those in focus in deeper planes were scored as transmigrated (brown arrows). **(J)** Quantitation of total adhered (surface adherent + transmigrated) and transmigrated neutrophils in TFM or TCM-MDA treated monolayers without TNFα treatment. Each data point represents the mean count of at least 12 high power fields of view (FOV). Unpaired t-test.

### Endothelial ACKR1 regulates neutrophil recruitment to the lung metastatic niche

Neutrophil recruitment creates a lung metastatic niche that supports tumor cell engraftment and proliferation. Neutrophils accumulate in the early lung metastatic niche at a similar stage of metastatic progression as seen for the upregulation of lung endothelial ACKR1^21,22,70,71^. Endothelial ACKR1 regulates neutrophil recruitment and extravasation in inflamed muscle tissue^42,43^. Therefore, we tested whether endothelial ACKR1 regulates neutrophil infiltration into the lung metastatic niche.

We quantified neutrophils in the same lung samples that we had examined for tumor cell extravasation, above. We found that Ackr1^ECKO^ mice had significantly fewer Gr-1^pos^ neutrophils in the lungs compared to controls and neutrophils comprised a smaller percentage of the immune infiltrate, indicating reduced neutrophil recruitment (Fig. 7D-F). We confirmed reduced neutrophil infiltration in Ackr1^ECKO^ lungs compared to Control using immunohistochemistry for Ly6G+ cells (Fig. 7G-H).

Loss of ACKR1 from RBCs can cause neutropenia^72,73^, and the observed reduction in neutrophil infiltration into the lung could be secondary to lower levels of circulating neutrophils. We measured circulating neutrophil levels in Control and Ackr1^ECKO^ mice before initiating TCM treatment and after tumor cell dissemination and found no indication of neutropenia in Ackr1^ECKO^ mice compared to Control mice (Fig. S4C). Together, these data indicate that endothelial ACKR1 does not affect the initial tumor cell extravasation into the lung parenchyma. Instead, endothelial ACKR1 plays a key role in recruiting neutrophils to the lung metastatic niche, which is known to support tumor cell engraftment and development of overt metastatic nodules.

### Tumor secreted factors promote endothelial ACKR1-mediated neutrophil adhesion and transmigration

We sought to determine how endothelial ACKR1 regulates neutrophil trafficking in a tumor-associated inflammatory setting. Previous work in inflammatory contexts has shown endothelial ACKR1 can localize CXCL1 and CXCL2 to the endothelial surface, promoting enhanced adhesion of neutrophils and extravasation of neutrophils, respectively^42,43^. We therefore hypothesized that endothelial ACKR1 might be enhancing neutrophil adhesion and/or transendothelial migration (TEM) in the metastatic niche, providing a mechanistic explanation for the reduction in neutrophils observed in the tumor-bearing Ackr1^ECKO^ mouse lungs.

To examine the effects of ACKR1 induction on neutrophil interactions with isolated endothelial cells, we induced ACKR1 in HUVEC by priming with TCM-MDA, as described in Fig. 3D-F. HUVEC monolayers were seeded on hydrated three-dimensional collagen gels and primed with either TFM or TCM-MDA for 24 hours. A subset of cultures was additionally activated with TNFα for 4 hours, which has previously been shown to induce neutrophil TEM^65,74^. Freshly isolated neutrophils were added to HUVEC monolayers and allowed to adhere and transmigrate for 6 minutes. Quantification showed a significant increase in both neutrophil adhesion and transmigration across TCM-MDA primed HUVEC monolayers compared to TFM primed HUVEC (Fig. 7I-J and Fig. S4D). Addition of TNFα increased the total number of adherent and migratory neutrophils but showed similar differences between TCM- and TFM-primed monolayers (Fig. S4E). Collectively, these results suggest that endothelial ACKR1 promotes metastasis by increasing neutrophil adhesion and extravasation in the metastatic niche.

## DISCUSSION

Despite significant improvements in the clinical care of breast cancer patients, metastasis remains the leading cause of breast cancer–related mortality¹. The clinical disparity is stark: while the five-year survival rate for patients with localized disease approaches 99%, it drops to under 30% for those presenting with distant metastases². This sharp contrast highlights that one major barrier to improving outcomes is the inability to regulate or inhibit the biological processes that enable disseminated tumor cells to colonize and proliferate within distant organs. Understanding how the metastatic niche forms and supports disseminated tumor cells is essential for developing therapeutic strategies to prevent metastatic disease.

In this study, we identify endothelial ACKR1 as a regulator of neutrophil recruitment to the lung metastatic niche and a vascular protein that promotes metastatic colonization within the lung parenchyma. Using both spontaneous and experimental metastasis models, we demonstrate that endothelial ACKR1 promotes metastatic colonization without altering primary tumor growth. Mechanistically, we demonstrate that endothelial ACKR1 is upregulated in the lung vasculature during breast cancer progression by factors secreted by the primary mammary tumor. ACKR1-positive venules serve as focal points for both tumor cell localization within the lung parenchyma and neutrophil accumulation. Endothelial-specific deletion of ACKR1 significantly reduces neutrophil recruitment and metastatic seeding of the lung. Our data indicate that endothelial ACKR1 functions early in the process of pre-metastatic niche formation, prior to arrival of metastatic cells. Together, these findings reveal a previously unrecognized pro-metastatic role for endothelial ACKR1 and suggest that tumor-derived signals drive an ACKR1-dependent vascular program that facilitates lung metastasis. The upstream tumor-derived factors responsible for ACKR1 induction remain incompletely defined. Prior reports and our own data implicate inflammatory cytokines such as TNF-α as candidate drivers, but further studies are needed to delineate the precise molecular pathways.

A central insight from our work is that endothelial ACKR1 expression in the lung vasculature is induced by tumor-secreted factors and restricted to venular endothelial cells. Upregulation of endothelial ACKR1 in the lung vasculature is observed in three separate experimental models, including the orthotopic breast cancer cell injection model, the MMTV-PyMT spontaneous breast cancer model, and with the administration of breast tumor-conditioned media (TCM) alone. Endothelial ACKR1 is typically restricted to venular endothelial cells, including HEVs and PCVs, where it regulates leukocyte extravasation^24,26,27,75-77^. Prior studies in lung allograft rejection, acute pulmonary infection, suppurative pneumonia, renal allograft rejection, rheumatoid arthritis, synovitis, and giant cell arteritis have demonstrated that endothelial ACKR1 is dynamically upregulated in response to inflammatory cytokines^29-31,33-35,78^. Our findings extend this paradigm to cancer metastasis by showing that breast tumors secrete factors that co-opt this inflammatory program to remodel ACKR1 expression in pulmonary venules and promote metastasis.

Multiple studies have demonstrated that ACKR1 expression is required for effective neutrophil recruitment to inflamed tissues^31,40,42,78^. However, earlier work has relied on global ACKR1 deletion in mouse models, leaving unresolved whether reduced neutrophil recruitment was a consequence of loss of endothelial ACKR1, which regulates chemokine presentation and transendothelial migration, or loss of erythrocyte ACKR1 expression, which controls circulating chemokine reservoirs^40,42,79^. Using an endothelial-specific *Ackr1* knockout model, we show that endothelial ACKR1 plays a significant role in neutrophil recruitment to the metastatic lung niche, as loss of endothelial ACKR1 results in a marked reduction in neutrophil accumulation. This Ackr1^flox^ mouse allele provides a powerful tool to investigate the functional role of endothelial *Ackr1* in breast cancer metastasis and other pathologies.

We demonstrate that endothelial ACKR1 deletion impairs lung metastatic growth. Reduction in neutrophil infiltration to the metastatic niche is likely to contribute to this reduction in metastasis, as neutrophils have an established role in pre-metastatic niche formation and support of metastatic tumor cell survival^17,21,22^. However, neutrophils constitute a heterogeneous and highly plastic population, and whether loss of specific neutrophil subsets contributes to diminished metastatic burden in our model remains unresolved. Notably, neutrophils can adopt distinct anti-tumorigenic (N1) or pro-tumorigenic (N2) states in response to tumor-derived cues, raising the possibility that endothelial ACKR1 preferentially supports recruitment of pro-tumorigenic neutrophils to the lung metastatic niche^18^. Defining the phenotypic and functional states of neutrophils recruited by ACKR1 expression will be an important direction for future studies. Further, ACKR1 has also been shown to facilitate extravasation of other cell types, such as monocytes and T cells^40,80^. Additional investigation is needed to determine whether ACKR1 directly influences recruitment, extravasation, and activation of other immune cell types relevant to metastatic progression.

Several complementary mechanisms may explain how endothelial ACKR1 regulates neutrophil recruitment in the metastatic lung. Our results show that tumor-conditioned media from both mouse and human breast cancer cell lines induce ACKR1 expression in endothelial cells, suggesting that tumor-derived inflammatory factors remodel venular endothelial signaling before the arrival of appreciable tumor cells. Prior mechanistic work demonstrates that ACKR1 binds CXCL1 on the endothelial surface to enhance neutrophil adhesion and concentrates CXCL2 at endothelial junctions to guide transendothelial migration^32,40,42,81^. Our *in vitro* studies support this model where tumor-conditioned endothelial monolayers exhibit increased neutrophil adhesion and transmigration. Coupled with our *in vivo* finding that Ackr1^ECKO^ mice maintain normal circulating neutrophil levels yet show reduced lung neutrophil infiltration, these results suggest that endothelial ACKR1 promotes neutrophil accumulation within the lung metastatic niche, potentially by facilitating neutrophil adhesion and transendothelial migration. Since neutrophils facilitate metastatic outgrowth through mechanisms that include suppressing anti-tumor immunity^18-22^, supporting tumor cell adhesion in the lung parenchyma through neutrophil extracellular trap secretion^17^, and promoting tumor cell extravasation across the endothelium^82^, ACKR1-dependent neutrophil recruitment may represent a key early event in establishing a metastasis-permissive vascular niche.

Although our data identify endothelial ACKR1 as an important regulator of lung neutrophil recruitment and metastatic outgrowth, several questions remain. First, as mentioned above, whether reduced lung metastasis in Ackr1^ECKO^ mice is driven primarily by impaired neutrophil recruitment or whether endothelial ACKR1 also influences tumor cell adhesion to endothelium, survival, or ability to extravasate remains to be determined. Second, the proximity of neutrophils and tumor cells to ACKR1-expressing vessels suggests that neutrophils and tumor cells preferentially extravasate from these venules, but we have recently shown that neutrophils extravasate preferentially across capillaries in the lung in several pathological contexts^66^. The lung metastatic niche may favor venular extravasation, as has been shown in other tissues^64,65^, or neutrophils and tumor cells may extravasate through capillaries and migrate into proximity of ACKR1-expressing venules due to favorable chemokine gradients or other microenvironmental cues. Third, while our endothelial-specific model isolates the vascular contribution of ACKR1, erythrocyte-specific ACKR1 is known to regulate circulating chemokine reservoirs and impact leukocyte recruitment to inflamed tissues^32,78,81,83-85^. Dissecting the relative contributions of endothelial versus erythrocyte ACKR1 pools will require the use of RBC-specific deletion models. Finally, determining the scope of endothelial ACKR1 function is critical for understanding broader applicability: further studies are required to determine whether endothelial ACKR1 is similarly upregulated in the vasculature of other metastatic target organs, whether differential ACKR1 induction guides metastatic “tropism” of different tumors, whether ACKR1 is upregulated by cancers other than mammary cancer, and whether endothelial ACKR1 is upregulated in the lung vasculature of human patients with breast cancer.

From a translational perspective, our work highlights endothelial ACKR1 as a potential therapeutic target for preventing or limiting breast cancer metastasis. Neutrophil infiltration is a consistent feature of metastatic niches, and elevated neutrophil-to-lymphocyte ratio is a well-established predictor of poor outcomes in patients with breast cancer and other malignancies^86^. By linking endothelial ACKR1 to neutrophil accumulation in the metastatic lung niche, our findings suggest that inhibition of this pathway could selectively disrupt pro-metastatic immune remodeling without broadly impairing systemic immunity. Moreover, endothelial ACKR1 induction may serve as a biomarker of pre-metastatic niche formation, with potential applications in risk stratification and early intervention.

Together, these findings position ACKR1 as a central regulator of tumor–endothelial–neutrophil interactions in the lung metastatic niche and establish upregulation of endothelial ACKR1 as an early event of the pre-metastatic niche. By using an endothelial-specific genetic model, we identify endothelial ACKR1 as a functional contributor to metastatic colonization of the lung, demonstrating that its loss reduces both neutrophil accumulation in the lung metastatic niche and metastatic burden. These insights define endothelial ACKR1 as a key vascular component influencing breast cancer metastasis, thereby expanding the framework through which endothelial responses are understood to shape metastatic progression. The discovery highlights multiple avenues for us to better understand the vascular programs that promote lung metastasis, and to consider therapeutic interventions based on this knowledge.

## MATERIALS AND METHODS

### Animal Models

All animal studies were approved by the Institutional Animal Care and Use Committee at the University of Illinois Chicago. All mice were maintained on a C57BL6/J background (Jackson Labs). Ackr1^Null^ mice^50^ were kindly provided by Dr. Victor Torres (New York University). *MMTV-PyMT* mice (Jackson Labs 022974) were provided by Dr. Pradip Raychaudhuri (University of Illinois Chicago). Routine genotyping of all alleles was performed by Transnetyx.

*Generation of Ackr1 Conditional mouse allele*: CRISPR-based Ackr1^flox^ mice were designed and generated in collaboration with Cyagen (Santa Clara, CA). Fertilized C57BL6/J mouse embryos were injected with guide RNA (gRNA) targeting exon2 of ACKR1, linearized repair template with loxP sites flanking exon 2, and Cas9 mRNA (Fig. 5A). Excision of Exon 2 via the resulting loxP sites results in removal of almost all of the *Ackr1* coding sequence. Four F1 offspring from a mosaic F0 animal were confirmed for germline transmission via PCR (primers: TTCCCTCCATCTATCTGCTCTCTTA and TAGATAGAAGGGGCATTTGGGTTT) and by sequencing the entire *Ackr1* locus amplified using primers external to the repair template.

*Ackr1 Endothelial Cell Knockout mice* (Ackr1^ECKO^). *Cdh5(PAC)-CreER^T2^* (*Cdh5CreERT2*)^68^ mice were bred into a background of homozygous *Ackr1^flox/flox^* mice to create mice with endothelial cell-specific deletion of *Ackr1*. Mice were bred to generate *Cdh5CreERT2;Ackr1^wt/wt^* (Control) and *Cdh5CreERT2;Ackr1^flox/flox^*homozygous (Ackr1^ECKO^) breeding cages for experimental cohorts. Cre-mediated gene recombination was induced with 100mg/kg tamoxifen (TAM; Sigma, T5648) dissolved in corn oil via oral gavage once daily for 5 days starting at 5 weeks of age.

### Cell Culture

E0771.LMB cells stably transfected with mCherry were provided by Dr. Andrew Dudley (University of Virginia)^52,87^. E0771.LMB cells were cultured in high glucose DMEM medium (Gibco, 11965092) supplemented with 10% FBS at 37°C with 5% CO_2_. The AT3 mouse mammary carcinoma cell line (Millipore Sigma, SCC178) was provided by Dr. Emrah Er (University of Illinois Chicago). AT3 cells were cultured in high glucose DMEM medium supplemented with 10% FBS, 2mM non-essential amino acids (Millipore Sigma, TMS-001-C), 15mM HEPES, and 1X β-mercaptoethanol (Millipore Sigma ES-007-E) at 37°C with 5% CO_2_. MDA-MB-231 cells were gifted from Dr. Alexandra Naba (University of Illinois Chicago) and maintained in high glucose DMEM medium supplemented with 10% FBS. Human Umbilical Vein Endothelial Cells from pooled donors (HUVECs, Promocell, C-12203) were cultured in Endothelial Growth Medium 2 (EGM, Promocell, C-22011) on Type I Collagen-coated plates at 37°C with 5% CO_2_. All cells were tested for mycoplasma before use in experimental studies (Mycoalert, Lonza, LT07-318).

### Tumor Modeling

#### Orthotopic Tumor Implantations

Female mice aged 8–10 weeks old were orthotopically implanted in the 4th mammary fat pad with 2x10⁵ mCherry-expressing E0771.LMB cells, with 4x10^5 mCherry-AT3 cells resuspended in 100uL DPBS (Gibco, 14190144), or with DPBS for mock implantation. Tumor growth was monitored by measuring tumor dimensions with calipers three times per week. Tumor volume was calculated using the formula: volume = (length x width²)/2. Mice were euthanized in cohorts at indicated timepoints during tumor progression or when the first mice of the cohort reached the humane endpoint (tumor diameter ≥ 2cm), which occurred at 28 days post-implantation for E0771.LMB cells and 40 days post-implantation for AT3.

#### MMTV-PyMT tumor model

Female *MMTV-PyMT* transgenic mice were monitored weekly until a palpable primary tumor was detected, then tumor volume(s) were measured with calipers three times per week. Mice were euthanized at the indicated time points or at humane endpoint (diameter of largest tumor ≥ 2cm).

### qPCR Analysis

Total RNA was isolated from homogenized tissue or cells using the RNeasy Mini Kit (QIAGEN, 74104), and cDNA was synthesized with the High-Capacity cDNA Reverse Transcription Kit (Applied Biosystems, 4368814) according to the manufacturer’s instructions. Quantitative real-time PCR (qRT-PCR) was performed on a ViiA 7 Real-Time PCR System (Applied Biosystems), in duplicates for each primer pair, using Fast SYBR Green Master Mix (Applied Biosystems, 4385612). Relative fold changes in mRNA expression were calculated using the 2^-ΔΔCt^ method, with expression levels normalized to β-actin.

Tumor Conditioned Media (TCM) Preparation and Injection

To prepare TCM for administration to mice, E0771.LMB or AT3 cells were plated on 100-mm tissue culture plates in their respective culture medium described above and grown to 60-70% confluence. Cells were washed twice in DPBS and then incubated in high glucose DMEM medium (Gibco, 11965092) without the addition of any supplements or FBS for 48 hours at 37°C and 5% CO_2_. TCM was collected with a 10-mL syringe, filtered through a 0.45μm filter, aliquoted, and immediately flash-frozen in liquid nitrogen. TCM was stored at -80°C until ready for use. Mice were given daily intraperitoneal injections of 300uL TCM for the number of days specified in each figure. To prepare TCM for administration to HUVEC, EGM2 was incubated with E0771.LMB or MDA-MB-231 cells for 48 hours, filtered and flash-frozen as above.

### Metastasis Modeling

#### Lung Metastasis from Orthotopic Mammary Implantation

Mice were euthanized at the indicated endpoint using isoflurane followed by cervical dislocation. Mice were perfused with 10 mL of sterile DPBS via the right ventricle, the lungs were inflated by intratracheal injection of 0.5–1 mL of 4% methanol-free paraformaldehyde (PFA, ThermoFisher, 043368.9M), and then lungs were fixed overnight in 4% methanol-free PFA at 4°C. The lungs were dehydrated in 15% sucrose for 6–8 hours, transferred to 30% sucrose overnight at 4°C, embedded in Tissue-Tek O.C.T. Compound (Sakura, 4583), and sectioned at a thickness of 10μm. Lung sections were thawed for 15 minutes at room temperature, washed three times in PBS, and mounted in VECTASHIELD PLUS Antifade Mounting Medium with DAPI (Vector Labs, H-2000). Tilescan images of whole lung sections were captured using 10x/0.32 NA objective using the Leica DMI8 inverted microscope. Higher-resolution images focusing on individual disseminated tumor cells were captured using 40x/0.60 NA objective. Lung metastases were quantitated using QuPath software. Individual cell outlines were generated using the QuPath cellular segmentation script by expanding each DAPI-labeled nucleus boundary by 5μm. Metastatic cells were identified as any cell outline containing mCherry signal. For larger metastatic nodules with groups of neighboring cells, each individual cell of the nodule was counted as an individual tumor cell. The metastatic burden of whole lung sections (i.e. tumor cells per mm²) was calculated at three distinct depths of each lung separated by at least 500 microns. These values were then averaged to determine the overall metastatic burden per mouse.

#### Experimentally Induced Metastasis

For experimental metastasis studies, mice received 8-9 days of intraperitoneal injections of 300uL tumor-conditioned media to prime the lung metastatic niche. Mice then received 3x10^5^ mCherry-E0771.LMB or mCherry-AT3 cells resuspended in 200uL sterile DPBS via tail vein injection. Two weeks after tumor cell injection, mice were euthanized by isoflurane followed by cervical dislocation and lungs harvested as above. Lungs were examined in whole mount immediately after initial inflation/fixation on an AxioZoom V16 microscope (Zeiss) at 7x/ magnification using a dsRed fluorescence filter to visualize mCherry signal. Metastases were quantified by direct visual inspection under the microscope on both the ventral and posterior lung surfaces, and the totals were summed per lung.

### Immunofluorescence and Immunohistochemistry Staining

Primary antibodies used included anti-mouse ACKR1 (1 μg/mL, a kind gift of Dr. Ulrich von Andrian, Harvard Medical School), anti-mouse CD31 (1:100, Cell Signaling, 77699), anti-mouse EphB4 (2.5 μg/mL, R&D Systems, AF446), anti-mouse Pan Cytokeratin (1:200, ThermoFisher/Bioss, BS-10403R), and anti-mouse Ly-6G (E6Z1T) (1:200, Cell Signaling, 87048). Secondary antibodies included anti-rat Alexa Fluor 488 and 647 (1:1000, Invitrogen, A48269 and A48272), anti-rabbit Alexa Fluor 555 and 647 (1:1000, Invitrogen, A32794 and A32795), anti-goat Alexa Fluor 488 and 555 (1:1000, Invitrogen, A-11055 and A-21432), biotinylated goat anti-rabbit secondary antibody (1:500, Vector Labs, BA-1000), and streptavidin-HRP (1:1000, ThermoFisher, 21126).

#### Immunofluorescence Staining

For immunofluorescent staining, lungs were inflated, fixed, dehydrated, and frozen in O.C.T. as described above. Tumors and lymph nodes were fresh-frozen in O.C.T. All tissues were sectioned at 10μm thickness. Tissue sections were thawed at room temperature for 15 minutes, washed three times with PBS, subjected to antigen retrieval in 6.0 sodium citrate solution by boiling in a microwave for 20 minutes, and then permeabilized with 0.2% Triton X-100 in PBS for 10 minutes. Blocking was performed using 10% Normal Donkey Serum in PBS-T (0.01% Tween in PBS) (Blocking solution) for 1 hour at room temperature. After blocking, sections were incubated with primary antibody cocktails overnight at 4°C in a humidity chamber. The next day, sections were washed three times in PBS-T, treated with secondary antibodies (1:1000 in blocking buffer) for 1 hour at room temperature, and washed again in PBS-T. Quenching was performed with the Vector TrueVIEW Autofluorescence Quenching Kit (Vector Labs, SP-8400-15) per manufacturer’s instructions, and sections were mounted in VECTASHIELD Vibrance Antifade Mounting Medium with DAPI (Vector Labs, H-1800). Images were captured as described above.

#### Immunohistochemistry Staining

Frozen section preparation was performed as described above. After permeabilization, sections were treated with BLOXALL Endogenous Blocking Solution (Vector Labs, SP-6000) and an avidin/biotin blocking kit (Vector Labs, SP-2001) according to the manufacturer’s instructions, then incubated in blocking solution for 1 hour at room temperature. After blocking, slides were incubated overnight at 4°C with primary antibody. The next day, sections were treated with biotinylated anti-rabbit secondary antibody for 30 minutes at room temperature and streptavidin-HRP for 30 minutes and washed in PBS. Signal was visualized using IMPACT DAB Substrate Kit, HRP (Vector Labs, SK-4105) for 10 minutes, after which sections were washed with water, dehydrated in increasing concentrations of ethanol, and mounted with Permount Mounting Media (Fisher Scientific, SP15). Imaging was performed with a Zeiss Axio Imager at 10x magnification.

### Image Analysis

#### Feature quantification

To quantify ACKR1-positive vessels, we performed an immunofluorescent stain for CD31, EphB4, and ACKR1, as described above. Pulmonary venules were identified by colocalization of CD31 and EphB4 signal. Venules were classified as positive if any CD31 and ACKR1 co-localization was observed. The entire tiled image for each lung section was examined. The area of lung tissue was computed using the annotation function in QuPath software. The total number of ACKR1-positive pulmonary venules per area (mm^2^) in each lung section was calculated to measure ACKR1-positive vessel density per lung and density was averaged across all sections per lung.

To quantify the number of neutrophils per lung, we identified neutrophils based on Ly6G-positive staining and quantified the number of Ly6G-positive cells per area (mm^2^) per lung section like above.

#### Proximity of tumor cells and neutrophils to ACKR1-positive or -negative pulmonary venules

Lung sections were stained for ACKR1, CD31, EphB4, and either Pan-CK (to label tumor cells) or Ly6G (to label neutrophils) as described above. Using QuPath analysis software, a semi-automated workflow was developed to measure the distance between the borders of pulmonary venules and nearby tumor cells or neutrophils. Pulmonary venules were identified by the colocalization of CD31 and EphB4 signals, while individual tumor cells and neutrophils (i.e. target cells) were identified through DAPI-based cellular segmentation and colocalization of Pan-CK or Ly6G signal within cellular outlines. A custom QuPath script was then applied to calculate the minimum distance between the venule wall and each target cell within a 150-micron radius. Target cells were binned into categories of 0-50, 51-100, or 101-150 microns to the nearest venule. The ACKR1 channel was turned off during analysis, and venules were categorized as ACKR1-positive or -negative after measurements were complete to ensure blind analysis. The custom QuPath script used for this analysis is available upon request.

### Tumor Cell and Neutrophil Extravasation

For tumor cell extravasation studies, mice received 8-9 days of intraperitoneal injections of 300uL tumor-conditioned media to prime the lung metastatic niche. 1x10^6^ mCherry-E0771.LMB cells were resuspended in 200 µL of sterile PBS and injected into the retro-orbital sinus of mice. After 24 hours, mice were intravenously injected with 3 µg of anti-mouse MHCI (H-2Kb) APC (eBioscience, 17-5958-82) to label intravascular tumor cells. The antibody was allowed to circulate for 3 minutes before the mice were euthanized using isoflurane followed by cervical dislocation.

Lungs were minced on ice using scissors and resuspended in dissociation buffer consisting of 20 mg/mL Type I Collagenase (Worthington, LS004217) and 0.1 mg/mL DNase I (StemCell, 7900) in Hank’s Balanced Salt Solution (with calcium and magnesium; Life Technologies, 14025-092). The tissue was incubated at 37 °C for two 15-minute intervals and homogenized through an 18-G needle 30 times after each incubation. Digested tissue was passed through a 40-µm cell strainer (Fisher Scientific, 08-771-7), washed with DPBS, and incubated in 3 mL of 1x RBC lysis buffer (BioLegend, 420301) on ice for 5 minutes. The lysis reaction was stopped by adding 10 mL of PBS. Cells were washed twice with DPBS and used for subsequent flow cytometry analysis.

### Flow Cytometry Analysis

Single-cell suspensions were incubated with Zombie Aqua Fixable Viability Kit (1:1000, BioLegend, 423101) in PBS at room temperature for 15 minutes in the dark. The cells were then washed with staining buffer consisting of 1% bovine serum albumin (Sigma-Aldrich, A9418) in PBS. The cells were blocked with 1 µg/100 µL TruStain FcX Plus (anti-mouse CD16/32) (Biolegend, 156604) for 10 minutes on ice, protected from light. Primary antibodies were added, and the cells were incubated for 30 minutes at 4°C in the dark. To examine extravasation of tumor cells, lung samples were incubated with anti-CD31 FITC (1:200, Biolegend, 102405) and anti-CD326 (1:50, Milltenyi Biotec, 130-102-927). To determine the number of neutrophils, lung samples were incubated with anti-CD45 FITC (1:800, Biolegend, 103107) and anti-Ly6G/C AF700 (1:100, eBioscience, 59-5931-82). To examine ACKR1 induction in HUVECs, samples were incubated with anti-CD31-FITC (1:80, BioLegend, 303103) and anti-ACKR1-PE (1:20, VWR, 35837). Following staining, the cells were washed twice with staining buffer and stored at 4°C until downstream flow cytometry analysis was performed.

For blood cell analysis, whole blood was collected via the facial vein into EDTA-lined BD Microtainer Capillary Blood Collector tubes (BD, 365974) and 100uL of whole blood was used for antibody staining. Primary antibodies were added to each sample and incubated for 30 minutes at 4°C, protected from light. For red blood cell ACKR1 expression studies, whole blood was incubated with anti-TER119 APC (1:200, Biolegend, 116211) and anti-ACKR1 AF647 (1:100, a kind gift of Dr. Ulrich Von Adrian) antibodies. Samples were washed twice with staining buffer and stored at 4°C for flow cytometry analysis.

All samples were gated on viable cells, followed by the exclusion of cell doublets. For neutrophil population studies, samples were gated first on the CD45+ population, followed by gating on Ly6G/C+ positive cells. The percentage of LY6G/C+ cells per total CD45+ population is reported. For red blood cell ACKR1 expression analysis, samples were gated on the TER-119+ population and then ACKR1+ cells. The percentage of ACKR1+ per total TER119+ cells is reported. Flow cytometry samples were recorded on CytoFLEX S Flow Cytometer (Beckman Coulter), and flow data was analyzed using Kaluza software (Beckman Coulter, v2.2).

To distinguish intravascular from extravascular tumor cells, mCherry-E0771.LMB tumor cells were identified within the lung cell population by gating for the mCherry-positive signal. The signal from the I.V. injected anti-MHC1 was then used to determine whether tumor cells were intravascular or extravascular. As negative controls to set background cutoff levels, mice were injected with tumor cells on day 9, but not injected with intravascular anti-MHC1 antibody. Tumor cells from whole blood collected at the same time as lung harvest were used as positive controls for successful MHCI staining. Samples with faint MHC1 signal were considered failed injections and excluded. The percentage of tumor cells that were extravascular (i.e. mCherry^pos^, MHC1^neg^) was quantified.

### Polymorphonuclear Neutrophil (PMN) Collection and Transmigration Assay

PMN transmigration assays were performed as previously described^74,88^. Briefly, PMNs were isolated from freshly obtained blood anticoagulated with sodium EDTA using PolymorphPrep (Cardinal Medical Associates). Isolated PMNs were resuspended in M199 medium containing 0.1% human serum albumen to a concentration of 5 × 10^5^ cells/ml.

Endothelial cell (EC) monolayers were grown on hydrogel and either left unstimulated or activated with TNF-alpha (25 ng/mL) for 4 hours prior to the assay. Before the addition of PMNs, EC monolayers were washed with warm M199. PMNs were then seeded on top of the EC monolayers and allowed to transmigrate for 6 minutes at 37°C. After incubation, non-adherent PMNs were removed by sequential washes with 1 mM EDTA in divalent cation free PBS, PBS with calcium and magnesium, then fixed in 2.5% glutaraldehyde (Electron Microscopy Sciences) in 0.1 M sodium cacodylate buffer (pH 7.4) overnight at 4°C, and stained with Wright-Giemsa. The hydrogel plug containing the EC-PMN co-culture was removed from the well and mounted between a glass slide and coverslip. Adherent and transmigrated neutrophils were visualized using a Zeiss Axio Imager using a 63x/1.4 oil objective. Adherent and transmigrated neutrophils were scored manually. In each field, neutrophils that had nuclei that were in sharp focus in the same plane as endothelial nuclei were scored as adherent, while neutrophils that had nuclei in sharp focus in the planes where collagen fibrils were evident were scored as transmigrated. For each condition, neutrophil counts were assessed until either 100 cells were observed or 25 high-power fields were examined across three replicate co-cultures. To quantify adhesion, the total number of neutrophils (adherent and transmigrated) per high-power field was averaged across wells. The percentage of transmigrated neutrophils (%TEM) was calculated as the ratio of transmigrated neutrophils to the total number of adhered and transmigrated neutrophils per well, averaged across replicates.

### qPCR Analysis

Total RNA was isolated from cells using the RNeasy Mini Kit (QIAGEN, 74104), and cDNA was synthesized with the High-Capacity cDNA Reverse Transcription Kit (Applied Biosystems, 4368814) according to the manufacturer’s instructions. Quantitative real-time PCR (qRT-PCR) was performed on a ViiA 7 Real-Time PCR System (Applied Biosystems), in duplicates for each primer pair, using Fast SYBR Green Master Mix (Applied Biosystems, 4385612). mRNA expression was normalized to β-actin, and relative expression levels were calculated using the 2^-ΔΔCt^ method.

### Statistics

All statistical analyses were conducted using GraphPad Prism 10.4.1 and the test used for each analysis is indicated in the relevant figure legend. Multiple tests corrections were used where indicated, and p values or corrected p values of less than 0.05 were considered significant. In graphs, a single data point represents a single animal and is the averaged value from a minimum of 3 quantified sections unless otherwise indicated.

### Supplemental material

Fig. S1 shows ACKR1 expression in primary tumors and tumor endothelium and the growth curves of primary tumors. Fig. S2 shows widespread induction of ACKR1 in mouse lungs by implanted mammary tumors, MMTV-PyMT mammary tumors, and IP-injected tumor-conditioned media. Fig. S3 shows co-localization of ACKR1 with EphB4 and detailed quantification of co-localization between ACKR1 and EphB4, ACKR1-expressing venules and tumor cells, and ACKR1-expressing venules and neutrophils. Fig. S4 shows control experiments for use of intravenous antibody to detect tumor cell extravasation, circulating neutrophil levels in Control and ECKO mice, and additional details of the neutrophil TEM assay.

## ACKNOWLEDGEMENTS

We would like to acknowledge the following funding sources: NCI predoctoral fellowship 1F30CA268764 (S. T. R.), American Heart Association predoctoral fellowship 25PRE1373228 (Q. W.), American Cancer Society I-CHER postdoctoral fellowship (C. E.), and American Cancer Society I-CHER/UI Cancer Center Pilot funding (C. E. and L. A. N.). We would like to thank the University of Illinois Cardiovascular Research Core and Jiwang Chen for assisting with tail vein injections and intratracheal TNF injections. We would like to thank the University of Illinois Chicago Flow Cytometry Core and Dr. Balaji Ganesh for assisting with flow cytometry design, optimization, and analysis. We would like to thank Ms. Lily Miller (Muller laboratory, Northwestern University) for assistance with HUVEC cultures for *in vitro* transendothelial migration experiments.

## Supplemental Figures

**Figure S1.**
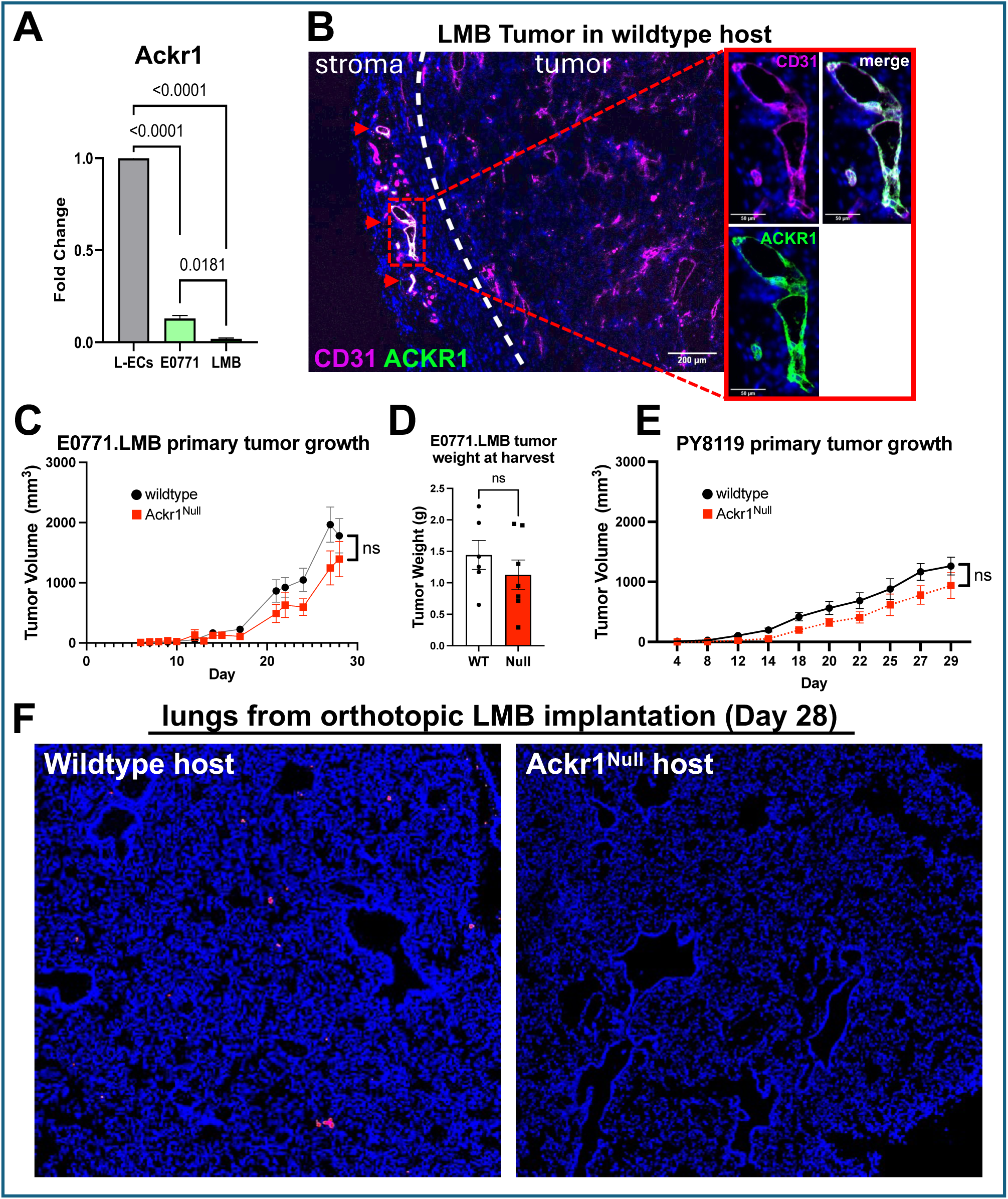
Host ACKR1 expression shows minimal effect on primary breast cancer growth. **(A)** Expression of ACKR1 mRNA in mouse lung endothelial cells (L-ECs), E0771 and E0771.LMB (LMB) cells. Significance determined by one-way ANOVA. **(B)** Representative image of Day 28 LMB tumor in a wildtype mouse, showing adjacent mammary stroma. Dotted line indicates border between tumor (right side of image) and stroma (left side of image). Rectangle indicates inset area adjacent. CD31 = magenta, ACKR1 = green. ACKR1 is expressed in some stromal vessels (red arrowheads). **(C-E)** Tumor growth curve and final weights of LMB and PY8119 tumors implanted into the 4^th^ mammary fat pad of WT and Ackr1^Null^ mice. Significance determined by multiple unpaired t-test (C, E) or unpaired t-test (D). **(F)** Representative low magnification overviews of lung sections of wildtype and Ackr1^Null^ mice implanted with LMB tumors, Day 28 endpoint. Red = RFP^pos^ tumor cells. Same mice as details shown in Figure 1A.

**Figure S2.**
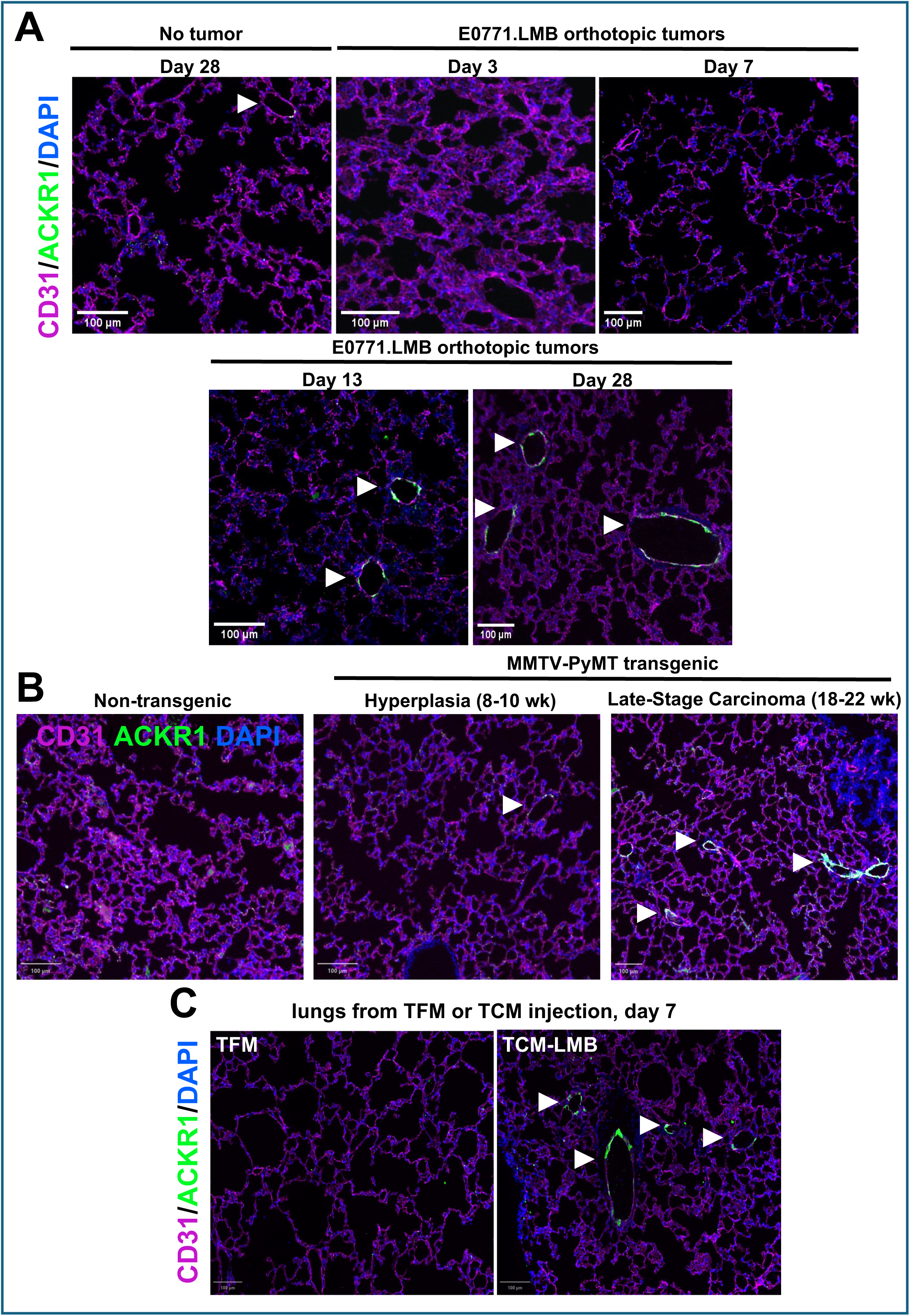
Induction of lung endothelial ACKR1 by implanted tumors, genetically induced mammary tumors, and secreted factors from tumors (low magnification overviews). **(A)** Representative low-magnification images of lungs from the indicated number of days after LMB implantation, showing the overall distribution of ACKR1 expression. **(B)** Representative low-magnification images of lungs from MMTV-PyMT mice at the indicated age showing the overall distribution of ACKR1 expression. **(C)** Representative low-magnification images of lungs from mice treated with TFM or TCM-LMB for 7 days, showing the overall distribution of ACKR1 expression. White arrowheads indicate ACKR1^pos^ vessels in all images.

**Figure S3.**
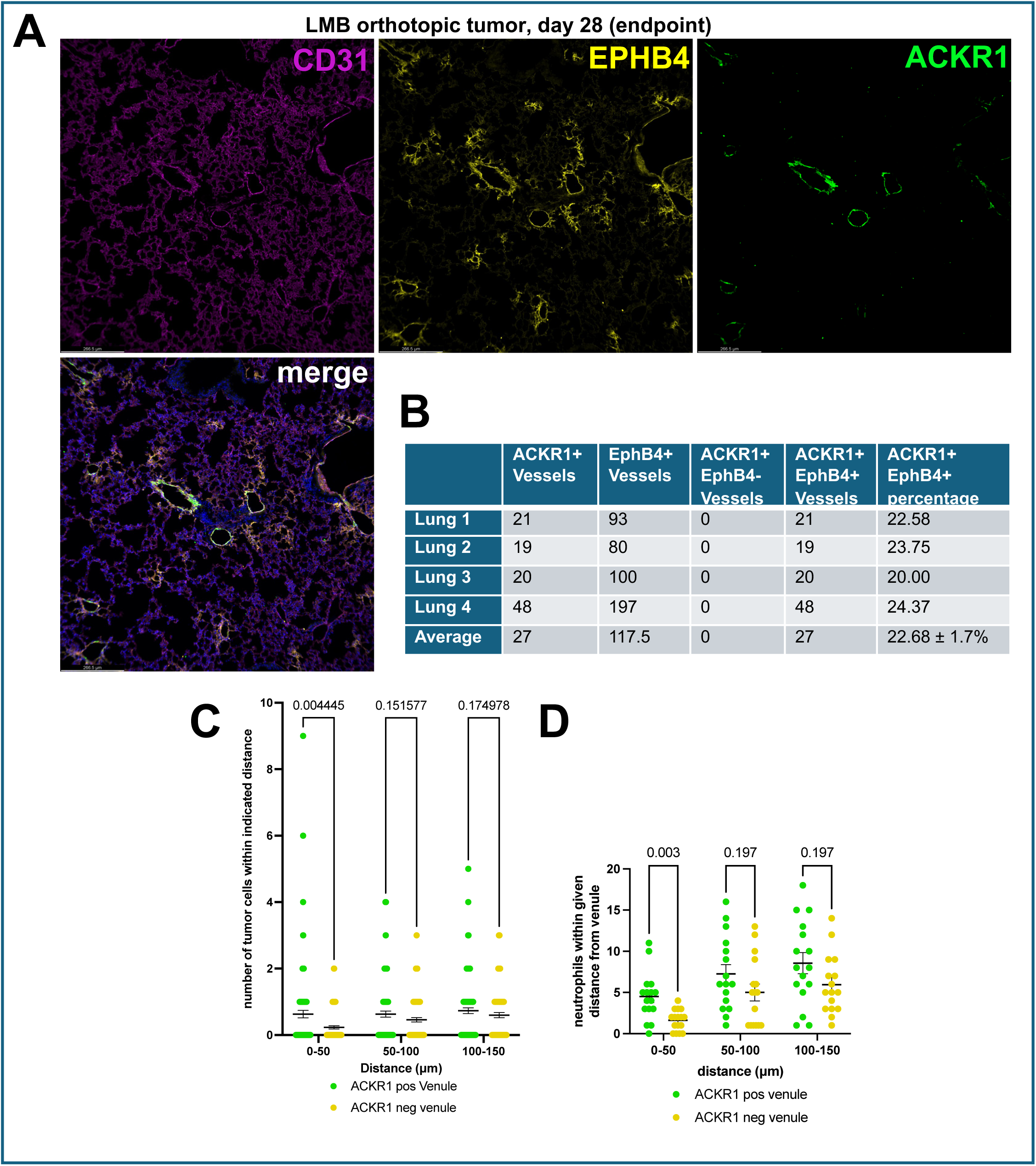
ACKR1 is upregulated specifically in a subset of venules. **(A)** Representative low-magnification images of lung from mice implanted with LMB tumors at Day 28, showing overall distribution of ACKR1 expression relative to EphB4 expression. **(B)** Quantification of vessels positive for ACKR1, EphB4, and both ACKR1/EphB4, across 4 individual mice. **(C)** Quantification of the number of DTCs within indicated distance from the border of ACKR1-positive or ACKR1-negative venules in Day 28 LMB-bearing mouse lungs (n = 116 ACKR1^pos^ venules, n = 100 ACKR1^neg^ venules, across 4 individual mice). Unpaired t-tests. Individual data points from graphs in Figure 4C. **(D)** Quantification of the number of neutrophils within indicated distance from the border of ACKR1-positive or ACKR1-negative venules in Day 28 LMB-bearing mouse lungs (n = 16 ACKR1^pos^ venules, n = 16 ACKR1^neg^ venules, across 3 individual mice). Unpaired t-tests. Individual data points from graphs shown in Figure 4F.

**Figure S4.**
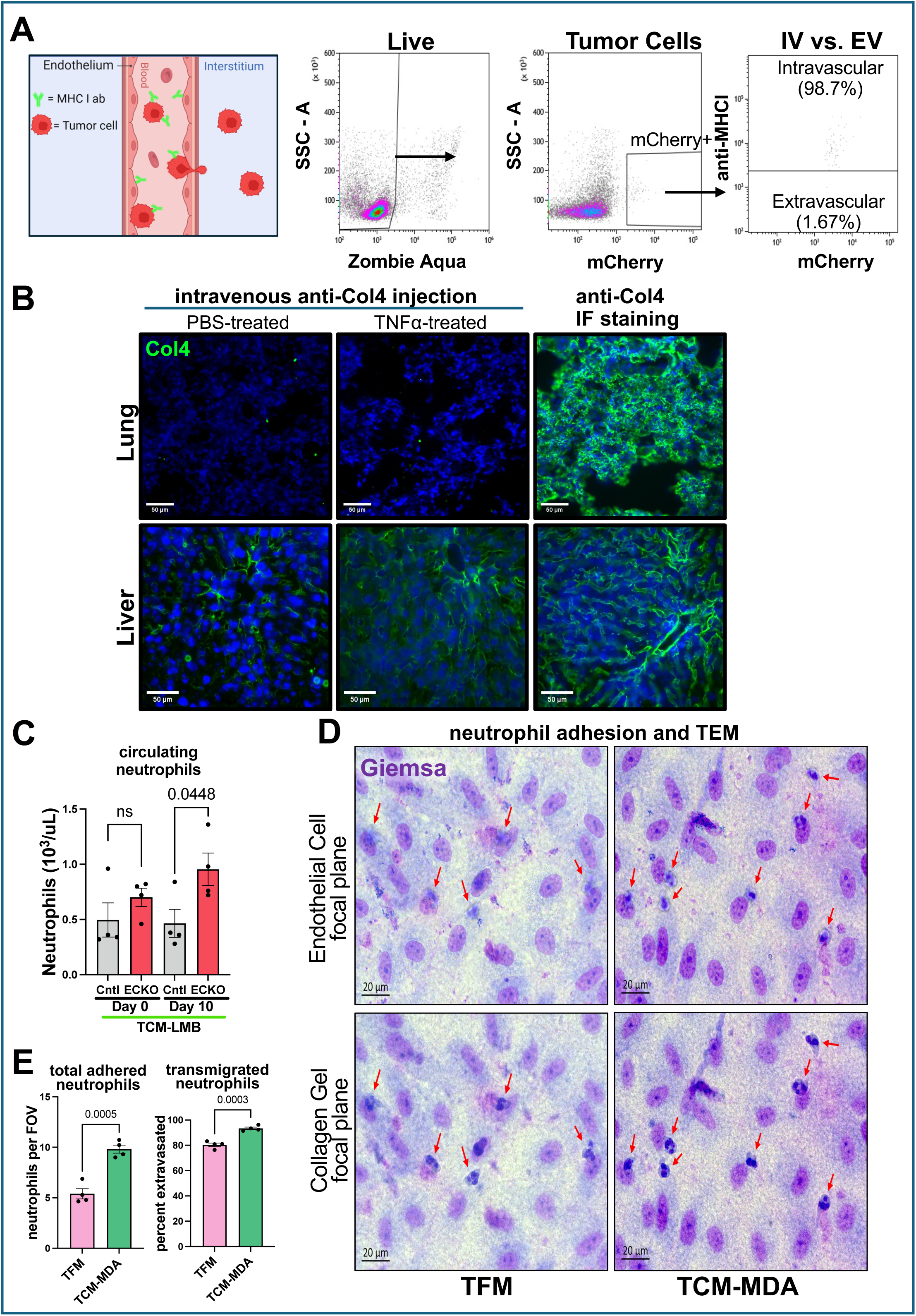
ACKR1 promotes neutrophil transendothelial migration. **(A)** Confirmation of MHCI labeling of intravascular tumor cells. Flow analysis of blood from animals in Figure 7A-C, showed that 99% of intravascular tumor cells were positive for anti-MHCI. **(B)** To examine antibody leakage to the extravascular space in the lung, we intravenously injected an antibody to collagen type IV (Col4), which does not have a target when confined to the endothelial lumen but readily binds the abluminal endothelial basement membrane upon leakage^73^. Mice were intratracheally injected with PBS (control) or 300 ng TNFα to induce lung inflammation. After 24 hours, anti-Col4 was administered via retroorbital injection three minutes prior to euthanasia. No Col4 leakage was observed in baseline or inflamed lungs (top left and center). Fenestrated liver endothelium acted as a positive control for Col4 leakage (bottom left and center). IF staining on sections confirmed Col4 immunoreactivity in both tissues (top and bottom right). **(C)** Absolute neutrophil counts (ANC) from automated complete blood counts (CBCs) of mice treated with TFM or TCM-LMB for 9 days, then injected retroorbitally with LMB tumor cells 24 hours prior to sacrifice (see figure 7, panels A-F). Circulating neutrophil levels were not reduced in ECKO animals. **(D)** Representative images of neutrophil transendothelial migration assays with TFM- or TCM-MDA-treated HUVEC. Red arrowheads indicate neutrophils, which may be in focus or out of focus in each focal plane. **(E)** Quantitation of total adhered (surface adherent + transmigrated) and transmigrated neutrophils in monolayers treated with TFM or TCM-MDA for 24 hours and then activated with TNFα 25 ng/mL for 4 hours prior to seeding with neutrophils. Each data point represents the mean count of at least 12 high power fields of view (FOV). Unpaired t-test.

